# A statistical simulator scDesign for rational scRNA-seq experimental design

**DOI:** 10.1101/437095

**Authors:** Wei Vivian Li, Jingyi Jessica Li

## Abstract

**Motivation:** Single-cell RNA-sequencing (scRNA-seq) has revolutionized biological sciences by revealing genome-wide gene expression levels within individual cells. However, a critical challenge faced by researchers is how to optimize the choices of sequencing platforms, sequencing depths, and cell numbers in designing scRNA-seq experiments, so as to balance the exploration of the depth and breadth of transcriptome information.

**Results:** Here we present a flexible and robust simulator, scDesign, the first statistical framework for researchers to quantitatively assess practical scRNA-seq experimental design in the context of differential gene expression analysis. In addition to experimental design, scDesign also assists computational method development by generating high-quality synthetic scRNA-seq datasets under customized experimental settings. In an evaluation based on 17 cell types and six different protocols, scDesign outperformed four state-of-the-art scRNA-seq simulation methods and led to rational experimental design. In addition, scDesign demonstrates reproducibility across biological replicates and independent studies. We also discuss the performance of multiple differential expression and dimension reduction methods based on the protocol-dependent scRNA-seq data generated by scDesign. scDesign is expected to be an effective bioinformatic tool that assists rational scRNA-seq experiment design based on specific research goals and compares various scRNA-seq computational methods.

**Availability:** We have implemented our method in the R package scDesign, which is freely available at https://github.com/Vivianstats/scDesign.

## 1 Introduction

The emergence and rapid development of single-cell RNA sequencing (scRNA-seq) technologies offer unprecedented opportunities for investigating transcriptional mechanisms underlying biological and medical phenomena at the individual-cell resolution (Wagner et al., 2016; Haque et al., 2017). While bulk RNA sequencing has been widely used to capture the average transcriptome information in a batch of cells (Li and Li, 2018b), scRNA-seq allows the investigation of transcriptome variation across thousands to millions of cells. The scRNA-seq technologies have enabled researchers to investigate fundamental biomedical questions such as cellular composition of various tissues and cell types, cell differentiation trajectories, and spatial and temporal dynamics of single cells. Important discoveries have been made from scRNA-seq data and advanced our understanding of diseases such as neurological disorders (Skene et al., 2018) and tumorigenesis (Tirosh et al., 2016).

Since the first scRNA-seq study was published in 2009 (Tang et al., 2009), more than twenty scRNA-seq experimental protocols have been developed. An effective experimental design requires careful consideration of the target research question as well as the experimental budget, and a typical design in practice consists of two steps. First, researchers need to select a proper protocol among the available ones, and the primary consideration is the choice between a tag-based protocol that allows the integration of unique molecular identifiers (UMIs) (Kivioja et al., 2012) and a full-length protocol that captures full-length transcripts and allows the addition of the External RNA Control Consortium (ERCC) spike-ins (Bacher and Kendziorski, 2016). The tag-based protocols (e.g., Drop-seq (Macosko et al., 2015)) are usually used to obtain a broad but shallow view of the transcriptomes across many cells, while the full-length protocols (e.g., Smart-seq2 (Picelli et al., 2013)) provide a deeper account of the gene expression in fewer cells. For example, a study about gene expression dynamics during stem cell differentiation requires accurate gene expression measurements, so it should opt for a full-length protocol. In contrast, in a study aiming to identify a previously unknown cell phase during the differentiation, it is necessary to sequence a large number of cells using a tag-based protocol to capture the transient phases. In the second step, to optimize an experiment with a selected protocol and a fixed budget, researchers need to choose between exploring the depth or breadth of transcriptome information, which sums up to determining the appropriate number of cells to sequence.

However, in contrast to the classical experimental design (Quinn and Keough, 2002) guided by certain theoretical optimality (e.g., the maximum power of a statistical test), the scRNA-seq experimental design is impeded by various sources of data noises, making a reasonable theoretical analysis tremendously difficult (Pierson and Yau, 2015; Kolodziejczyk et al., 2015). Especially, scRNA-seq data are characterized by excess zeros resulted from *dropout* events, in which a gene is expressed in a cell but its mRNA transcripts are undetected. As a result, many commonly used statistical assumptions are not directly applicable to modeling scRNA-seq data. For example, Baran-Gale et al. proposed using a negative binomial model to estimate the number of cells to sequence, so that the resulting experiment is expected to capture at least a specified number of cells from the rarest cell type (Baran-Gale et al., 2017). However, the estimation accuracy depends on the idealized negative binomial model assumption, which real scRNA-seq data usually do not closely follow (Figure S1). There is also a theoretical investigation of the cell-depth trade-off based on the Poisson assumption of gene read counts and a specific list of genes of interests (Zhang et al., 2018). In contrast to model-based design approaches (Dumitrascu et al., 2018b), multiple scRNA-seq studies used descriptive statistics to provide qualitative guidance instead of well-defined optimization criteria for experimental design (Grün and van Oudenaarden, 2015; Rizzetto et al., 2017). However, because the descriptive statistics were proposed from different perspectives, their resulting experimental designs are difficult to unify to guide practices. For example, one study reported that the sensitivity of most protocols saturates at approximately one million reads per cell (Ziegenhain et al., 2017), while another study found that the saturation occurs at around 4.5 million reads per cell (Svensson et al., 2017). The reason for this discrepancy is that the two studies defined the sensitivity in different ways: the first study used the gene detection rate while the second study used the minimum number of input RNA molecules required for confidently detecting a spike-in control (Jiang et al., 2011).

In this paper, we propose a statistical simulator scDesign for optimizing scRNA-seq experimental design from the perspective of detecting differentially expressed (DE) genes between two biological conditions (determined before an experiment) or two cell states (inferred after an experiment), a major scRNA-seq data analysis task. Given a pre-defined significance level (e.g., a false discovery rate), the power of an scRNA-seq experiment for detecting DE genes is jointly determined by the sensitivity of detecting gene expression, the accuracy of measuring gene expression, and the number of cells sequenced for each cell state. For each protocol and a specified total sequencing depth (i.e., the total number of reads in an scRNA-seq experiment), the cell-wise sequencing depth (i.e., the expected number of reads per cell) decreases as the cell number increases (Haque et al., 2017). However, existing power analysis methods for scRNA-seq experiments unrealistically assume a fixed cell-wise sequencing depth, which does not change as the cell number varies (Ziegenhain et al., 2017; Vieth et al., 2017). Therefore, the practical scRNA-seq experimental design calls a new approach that accounts for various characteristics and constraints of a real scRNA-seq experiment.

ScDesign is a simulation-based experimental design framework that has multiple advantages in real practice. First, scDesign is protocol- and data-adaptive. It learns scRNA-seq data characteristics from rapidly accumulating public scRNA-seq data generated under diverse settings. For example, 1, 976 series of scRNA-seq datasets are currently available in the Gene Expression Omnibus (GEO) database (Edgar et al., 2002). There are also newly developed scRNA-seq databases such as SCPortalen (70 studies with 67, 146 cells) (Abugessaisa et al., 2017), scRNASeqDB (36 studies with 8, 910 cells) (Cao et al., 2017), and the Single Cell Portal (43 studies with 496, 366 cells). Second, scDesign generates synthetic data that well mimic real scRNA-seq data under the same experimental settings, providing a basis for using its synthetic data to guide practical scRNA-seq experimental design. Third, scDesign is flexible in accommodating user-specific analysis needs. Users can apply scDesign to evaluate the performance of downstream analysis, such as gene differential expression and cell clustering, under various experimental settings at no experimental cost. Assisted by the evaluation results, users will be able to design an scRNA-seq experiment based on the setting leading to the best performance according to their specified criteria.

## 2 Methods

### 2.1 The statistical framework of scDesign

We develop scDesign based on a statistical generative framework that utilizes both existing real scRNA-seq data and reasonable assumptions mimicking various experimental processes. In contrast to the existing simulation methods for scRNA-seq data, scDesign constructs a Gamma-Normal mixture model to account for dropout events. This is motivated by the successful applications of our previously developed imputation method, scImpute, for recovering dropout gene expression values in scRNA-seq data (Li and Li, 2018a). This mixture model allows scDesign to overcome the dropout hurdle in learning the key gene expression characteristics from real scRNA-seq data (Figure S1), so that scDesign generates synthetic data highly similar to real data in multiple aspects. Depending on whether the task is to design an scRNA-seq experiment to sequence one or two batches of cells, scDesign has the corresponding one-state mode (Figure S2a) or the two-state mode (Figure S2b). In the one-state mode, scDesign leverages the information in a real scRNA-seq dataset from one biological condition (e.g., treatment or control) or one cell state (e.g., T cells) to generate a single scRNA-seq dataset given an experimental setting, i.e., a pre-specified total sequencing depth and a cell number. From the real scRNA-seq dataset, scDesign first estimates two cell-wise and three gene-wise parameters, which jointly define the key characteristics of scRNA-seq data. Second, scDesign simulates ideal gene expression levels for new cells of the same biological condition or cell state based on the estimated gene expression parameters. Third, scDesign introduces missing values to mimic the actual dropout events in an scRNA-seq experiment. Fourth, scDesign outputs a synthetic gene expression matrix with entries as read counts. In the two-state mode, scDesign leverages the information in two real scRNA-seq datasets from different biological conditions or cell states to generate two scRNA-seq datasets given an experimental setting. In this two-state mode, the simulation by scDesign mimics an experiment where two groups of cells from two biological conditions or cell states are sequenced together (Figure S2b).

### 2.2 scDesign for scRNA-seq data simulation

In this section, we describe how scDesign generates simulated RNA-seq data given existing real scRNA-seq data from a certain cell state. These simulated count matrices capture the characteristics of real count matrices, and they thus can be used to assist the development of computational methods and evaluate the performance of those methods under user-specified settings. We introduce how to simulate a single count matrix below, and please refer to the Supplementary Information for simulating multiple count matrices following a differentiation path.

Given a real single-cell count matrix with *I* genes and *J*_0_ cells, our goal is to generate a new count matrix with *I* genes and *J* cells, under the constraint that the new matrix has a total of *S* reads (Figure S2a). Both *J* and *S* are user-specified parameters. This resembles the real scenario where both the cell number and the total read number (i.e., the total sequencing depth) need to be specified before an scRNA-seq experiment.

#### 1. Estimate parameters from real scRNA-seq data

Denote the real single-cell count matrix by ***X***^real^, whose rows *I* and *J*_0_ columns represent the genes and cells, respectively. About the two cell-wise parameters, for each cell *j* we estimated its library size as

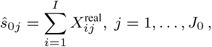

and its cell-wise dropout rate as

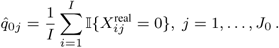

Then we fit the cell library sizes *ŝ*_01_, …, *ŝ*_0_*J*_0_ using a truncated Normal distribution, and the estimated mean and standard deviation (sd) are denoted as 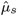 and 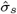, respectively.

To estimate the three gene-wise parameters, we first normalized the read counts given their corresponding library sizes and then performed a logarithmic transformation on the normalized values. The transformed matrix is denoted as ***X***^log^, where

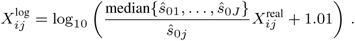

Using the Gamma-Normal mixture model described in the scImpute method (Li and Li, 2018a), we estimated the gene-wise dropout rate and mean and sd of gene expression. The mixture model consdiers the expression levels of gene *i* as independently and identically distributed random variable 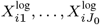 following the density function

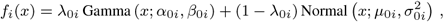

where λ_0*i*_ is gene *i*’s dropout rate, *α*_0*i*_ and *β*_0*i*_ are the shape and rate parameters of the Gamma distribution, and *μ*_0*i*_ and *σ*_0*i*_ are the mean and sd of the Normal distribution. The Gamma component describes the gene expression distribution when dropout occurs, while the Normal component represents the distribution of actual gene expression levels. We use multiple real scRNA-seq datasets to demonstrate that this mixture model outperforms the widely used Negative Binomial model in terms of goodness-of-fit to real data (Figure S1). The parameters in this model can be estimated by the Expectation-Maximization (EM) algorithm and the resulting dropout rate, mean, and sd estimates are denoted as 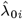, 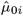, and 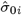, respectively. We then used a Gamma distribution to fit the estimated gene mean expression levels 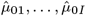 and denoted the estimated shape and scale parameters as 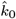 and 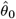.

To summarize, we estimated two cell-wise parameters including the cell library size *ŝ*_0*j*_ and the cell-wise dropout rate 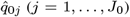, and estimated three gene-wise parameters including the mean expression 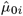, the sd 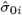, and the gene-wise dropout rate 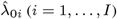.

#### 2. Simulate ideal gene expression values

In this step, we simulated the ideal expression values independently for each gene without considering varying cell library sizes and the dropout issue. For each gene *i* (*i* = 1,…,*I*), we first simulated its mean expression 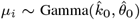, then we simulated its sd by stratified sampling from the binned observations, which we processed from the real count matrix. Specifically, we divided the estimated gene mean expression values 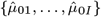 into *B* intervals, and we used 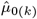 to denote the *k*-th order statistic of 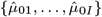. Then, the first interval was 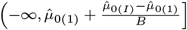, the *b*-th interval (1 < *b* < *B*) was 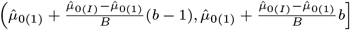, and the *B*-th interval was 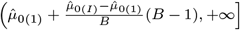. We defined 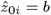 if 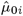 belonged to the *b*-th bin, and similarly we defined *z*_*i*_ = *b* if *µ*_*i*_ belonged to the *b*-th bin. We simulated the sd *σ*_*i*_ of gene *i* by sampling from the stratified gene sds estimated from the real data: 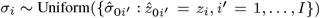. Finally, we generated the ideal expression matrix ***X***^ideal^, where 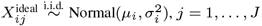.

#### 3. Introduce dropout events

In this step, we introduced dropout events into the synthetic count matrix, while accounting for the variability of both gene-wise and cell-wise dropout rates. The cell-wise dropout rate in a synthetic cell *j* was simulated as 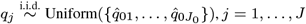, *j* = 1,…, *J*. For each gene *i* (*i* = 1,…, *I*), we simulated its gene-wise dropout rate *λ*_*i*_ by sampling one value from the stratified dropout rates estimated from the real data: 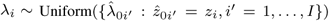. Then, we simulated the number of dropout events of gene *i*: *n*_*i*_ ~ Binomial(*J*,λ_*i*_). In other words, gene *i* was affected by the dropout events in *n*_*i*_ cells. These *n*_*i*_ cells were sampled without replacement from the cell population {1, 2,…,*J*, with cell *j* being selected with probability 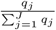. We denoted the sampling results by *I*_*ij*_, with *I*_*ij*_ = 1 indicating that gene *i* is a dropout in cell *j* and *I*_*ij*_ = 0 indicating that gene *i* is successfully amplified in cell *j*, *j* = 1,…,*J*. Then we obtained the synthetic count matrix with dropout events ***X***^drop^, where 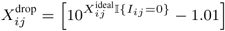, and [*x*] is the nearest integer to *x*.

#### 4. Simulate the final count matrix

We first simulated the library size of each synthetic cell 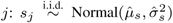, *j* = 1,…,*J*, and then we calculated the expected proportion of each entry in the count matrix

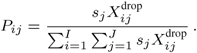

Finally, we obtained the final synthetic count matrix ***X***^syn^, which is constrained by the sequencing depth *S*, by simulating its counts from the multinomial distribution: 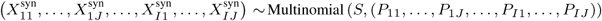.

### 2.3 scDesign for scRNA-seq experimental design

scDesign aims to determine the best number of cells to sequence given a fixed sequencing depth, such that the resulting RNA-seq data are optimized for differential gene expression analysis. We denote the two real count matrices as ***X***^real1^, with *I* rows (genes) and *J*_01_ columns (cells), and ***X***^real2^, with *I* rows (genes) and *J*_02_ columns (cells). Without loss of generality, we assume that the two matrices, which represent two cell states, have the same genes listed in the same order. We introduce how to simulate a synthetic count matrix for each state with scDesign in two scenarios, and the procedure is then repeated with varying cell numbers to obtain synthetic data for power analysis (Supplementary Information).

#### Scenario (1)

In scenario (1), we assume that cells from the two cell states are prepared as separate libraries and sequenced independently. Given ***X***^real1^ and ***X***^real2^, the goal of scDesign is to generate a synthetic count matrix with *I* genes and *J*_1_ cells for state 1, and a synthetic count matrix with *I* genes and *J*_2_ cells for state 2. Cell states 1 and 2 have sequencing depths of *S*_1_ and *S*_2_, respectively. For each state *g* (*g* = 1, 2), we followed Section 2.2 to simulate a count matrix 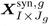. The only difference is in step 2, where we directly set 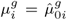 and 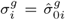, *i* = 1,…, *I*, instead of simulating new parameters. This is to ensure that the rows in the two simulated matrices still represent the same set of real genes, and the power analysis based on the simulated data is biologically meaningful.

#### Scenario (2)

Now we consider the case where the two cell states are jointly sequenced. Suppose that the two cell states are mixed in one biological sample, and the experimental setting is that *J* cells are to be sequenced to generate *S* RNA-seq reads in total. We assume that the two cell states present in fractions of *p*_1_ and *p*_2_ in the sample, respectively. That is, 0 < *p*_1_ < 1, 0 < *p*_2_ < 1, and *p*_1_ + *p*_2_ ≤ 1. When *p*_1_ + *p*_2_ < 1, there are more than two cell states present in the same sample. The goal of scDesign in scenario (2) is to simulate count matrices for the two selected cell states, based on a real count matrix of each state (Figure S2b).

1. Determine cell numbers We denote the numbers of cells from state 1, state 2, and the remaining states as *J*_1_, *J*_2_, and *J*_*r*_, respectively. These numbers were sampled from a Multinomial distribution: (*J*_1_, *J*_2_, *J*_*r*_) ~ Multinomial (*J*, (*p*_1_, *p*_2_, 1 − *p*_1_ − *p*_2_)).
2. Simulate count matrices with dropout events Following step 1-3 in Section 2.2, we simulated two count matrices 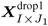 and 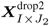 for cells states 1 and 2, respectively. The only difference was in step 2, where we directly set 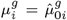 and 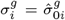, *i* = 1,…, *I*, to ensure that the rows in the synthetic count matrices represented the same set of real genes.
3. Simulate the final count matrices We first simulated the library sizes of the cells in the two states:

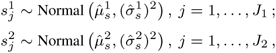

where 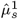 and 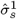 are estimated from ***X***^real1^, and 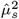 and 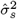 are estimated from ***X***^real2^. Then we combined the two count matrices to obtain the expected proportion matrix ***P***_*I*×(*J*_1_ + *J*_2_)_:

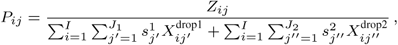

where 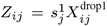 if 1 ≤ *j* ≤ *J*_1_, and 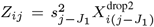 if *J*_1_ < *j* ≤ *J*_1_ + *J*_2_. The first *J*_1_ columns and the last *J*_2_ columns in ***P*** give the expected proportions of genes in cell states 1 and 2, respectively. We further assume that the total number of reads from the two states together is *S*_0_ = [*S*(*J*_1_ + *J*_2_)/*J*], where [*x*] denotes the nearest integer to *x*. Then we simulated the final count matrix 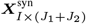 constrained by the sequencing depth from a Multinomial distribution: 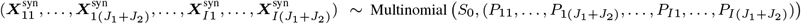. The final count matrix of cell state 1 and state 2 are 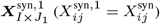 and 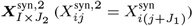, respectively.

## 3 Results

### 3.1 scDesign captures key characteristics of scRNA-seq data

We first demonstrate that scDesign accurately captures six key characteristics of real scRNA-seq data, so it serves as a reliable data simulator to assist scRNA-seq experimental design and to compare relevant computational methods. To assess the simulation performance of scDesign as compared with four other simulation methods, splat, powsimR, Lun, and scDD, we compared the simulated data generated by each method with the real data from various protocols. Both splat and powsimR are software packages for simulating scRNA-seq data (Zappia et al., 2017; Vieth et al., 2017); Lun denotes the simulation design introduced by Lun et al. (Lun et al., 2016); scDD denotes the simulation method designed to evaluate the DE method scDD (Korthauer et al., 2016). We considered six experimental protocols, Smart-seq2 (Picelli et al., 2013), Drop-seq (Macosko et al., 2015), 10× Genomics (Zheng et al., 2017), Fluidigm C1 (SMARTer) (Pollen et al., 2014), inDrop (Klein et al., 2015), and Seq-Well (Gierahn et al., 2017), and we collected three real scRNA-seq gene expression datasets of distinct cell types from each protocol (Table S1). In summary, we used 18 real count matrices of 17 cell types from human and mouse to evaluate the five simulation methods.

For each real count matrix, we randomly split the columns (cells) into two subsets of equal sizes, one used to estimate gene expression parameters and simulate a new count matrix with the same dimensions, and the other used to evaluate the simulation results. We compared each pair of real and simulated count matrices in terms of six summary statistics, including four gene-wise statistics (the count mean, the count variance, the count coefficient of variation (cv), and the gene-wise zero fraction) and two cell-wise statistics (the library size and the cell-wise zero fraction) (Supplementary Information). Our results show that scDesign well mimics real scRNA-seq experiments based on all the six experimental protocols, even though those protocols generate data with distinct properties. For example, data from Smart-seq2 and Fluidigm C1 have relatively larger library sizes and smaller count cvs (Figures 1a, S3, S4), while data from the other four protocols have smaller library sizes and larger zero proportions (Figures S5-S8). We measured the similarity between each summary statistics’ empirical distributions in real and the corresponding simulated data by each simulation method, using the Kolmogorov-Smirnov (KS) distance, whose value is between 0 and 1 and a smaller value indicates greater similarity (Supplementary Information). Comparing the KS distances of the five methods, we found that scDesign performs the best for five protocols: Smart-seq2, Fluidigm C1, Seq-Well, Drop-Seq, and inDrop (Figures 1b, S3, S4, S5, S7, S8), while scDesign and powsimR perform comparably for 10× Genomics (Figure S6). In summary, scDesign is ranked the best in 84 comparisons, among all the 108 comparisons (six statistics for each of the 18 datasets). In addition, our results also show that scDesign is able to preserve the relationships between genes’ expression mean and expression variance, expression cv, and zero proportion (Figure 1c). The demonstrated advantage of scDesign is rooted in its ability to incorporate both parametric and non-parametric methods to simulate scRNA-seq data. By constructing a mixture model to account for the dropout events, scDesign explicitly models the gene-wise parameters from the real data. When generating cellwise parameters for the simulated cells, scDesign uses different sampling techniques for each parameter to capture its distribution characteristic. In terms of the method stability, scDesign, Lun, and splat successfully estimated simulated data for all the 18 datasets, while scDD encountered errors for five datasets, and powsimR had errors with four dataset.

**Fig. 1.**
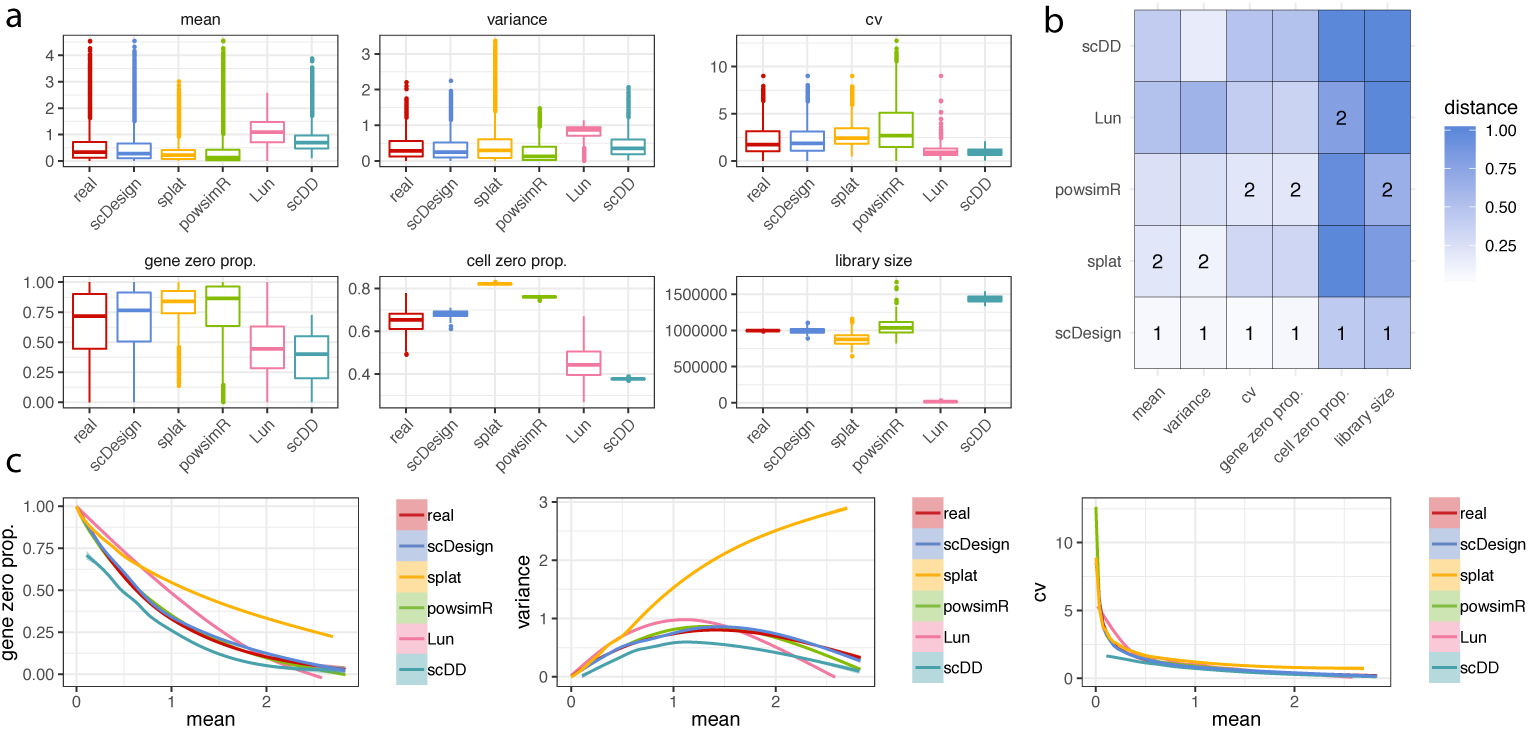
Comparison of scRNA-seq simulation methods based on the Smart-seq2 protocol. **a:** The gene-wise expression mean, expression variance, expression coefficient of variation, zero proportion, and the cell-wise zero proportion and library size in both real (monocytes) and simulated datasets. **b:** The KS distances between the six statistics in the real data and in the simulated data. The best and second best simulation methods with respect to each statistic are respectively marked with 1 and 2 in the heatmaps. **c:** The empirical relationships between the key statistics in the real and simulated data.

### 3.2 scDesign guides rational scRNA-seq experimental design

Given a fixed sequencing depth in designing an scRNA-seq experiment, scDesign assists users to predict the optimal number of cells for sequencing. In the context of gene differential expression analysis of two cell states, the cell number is optimal if its resulting scRNA-seq data lead to the most accurate detection of DE genes, where the accuracy depends on a user-specified criterion, e.g., a statistical test’s power given a significance level.We consider two scenarios: (1) cells from the two cell states are prepared as two separate libraries and sequenced independently; (2) cells from the two cell states are prepared in the same library and sequenced together. Scenario (1) includes many studies that investigated cells collected at two differentiating time points, cells of the same tissue type from patients and healthy subjects, or cells of the same type but exposed to different experimental treatments (Jaitin et al., 2014; Shekhar et al., 2016). The experimental design under scenario (1) aims to select the optimal cell numbers simultaneously for two libraries, so that the subsequent DE analysis becomes the most accurate given a user-specified criterion. Scenario (2) includes many scRNA-seq studies that sequenced an *in vivo* tissue sample, e.g., the peripheral blood mononuclear cell sample (Zheng et al., 2017), which is composed of a mixture of cell subtypes. In scenario (2), DE analysis is performed on a pair of known or putative cell subtypes within the sequenced sample. We consider the experimental design to optimize the DE analysis between two pre-selected cell subtypes under scenario (2).

In scenario (1), the constraints are the total sequencing depths of the two cell states, and scDesign aims to determine the optimal cell number for each cell state, among a set of candidate cell numbers. scDesign simulates a new count matrix of each state based on a real count matrix of the same state, for each pre-specified sequencing depth and cell number. Once obtaining the simulated count matrices corresponding to various candidate cell numbers, scDesign assesses the accuracy of DE gene identification using five metrics: precision, recall, true negative rate, F1 score (the harmonic mean of precision and recall), and F2 score (the harmonic mean of true negative rate and recall) (Table S2). We applied scDesign to optimize the designs of 14 example experiments (Table S3). In every experiment, we set the sequencing depth to 100 million reads, and considereded eight candidate cell numbers per cell state: 64, 128, 256, 512, 1024, 2048, 4096, and 8192. The DE genes between two cell states were identified using the two-sample *t* test.

Our results suggest that given a selected criterion in the DE analysis, the optimal cell number is jointly determined by multiple technical factors, including the experimental protocol and the variation introduced by sequencing, as well as biological factors, such as the intra- and inter-state cellular heterogeneity. Two factors are notable. First, when cells of the same two states are sequenced, the optimal cell number varies with protocols. For example, between two subtypes of glial cells: astrocytes and oligodendrocytes, 512 cells per state is the optimal cell number that maximizes the recall in DE analysis when Fluidigm C1 is used, but the number becomes 4096 per state when inDrop is used (Figure 2). If users choose the F1 score as the criterion, the optimal cell number per state is 128 and 1024 for Fluidigm C1 and inDrop, respectively. Therefore, Fluidigm C1 and inDrop require vastly different cell numbers to reach the same level of accuracy in DE analysis, and inDrop generally needs more cells than Fluidigm C1. This result is reasonable, since inDrop is a tag-based protocol that is advantageous in capturing more cells but disadvantageous in measuring each cell accurately, compared with the full-length protocol Fluidigm C1. Second, under the same protocol, the optimal cell number depends on the transcriptome similarity of the two cell states. For instance, with Smart-seq2, 512 cells need to be sequenced per state to maximize the recall in identifying DE genes between two dendrocyte subtypes, but only 256 cells per state are needed when dendrocytes are compared with monocytes (Figure S9). If the goal is to maximize the F2 score, the optimal cell number for comparing the two dendrocyte subtypes remains 512 per state, but the number reduces to 128 for comparing dendrocytes with monocytes. It is worth noting that the optimal cell number for both comparisons becomes 64, the smallest candidate cell number, when the criterion is the precision or the true negative rate (Table S3). The reason is that only the genes with strong DE signals are detectable with a small sample size (cell number) in any statistical testing. Hence, with a reasonable lower bound on the cell number, the DE genes detected at a smaller cell number have a higher precision. Unlike the precision, the largest recall in DE analysis is mostly achieved at a medium to large cell number. In all the experimental designs we evaluated, the recall rate of DE genes first increases with the cell number and then decreases after reaching a peak (Figures 2 and S9, Table S3). These results demonstrate the trade-off between the cell number and the cell-wise library size in scRNA-seq experiments. A combination of a small cell number and a large cell-wise library size ensures the identification of the DE genes with strong DE signals (i.e., achieving a high precision rate), but the small cell number may prohibit the detection of the DE genes with small to medium DE signals (i.e., sacrificing the recall rate). On the other hand, a combination of a reasonably large cell number and a small cell-wise library size increases the recall rate in detecting DE genes but compromises the precision rate due to high dropout rates (Figure S10). We also performed the DE analysis by replacing the two-sample *t* test with an scRNA-seq DE method MAST (Finak et al., 2015) (Table S4). The optimal cell number remains 64 per state when the criterion is the precision. The optimal cell numbers defined by the recall have small differences from the *t* test results, but the scale and trend remain largely consistent.

**Fig. 2.**
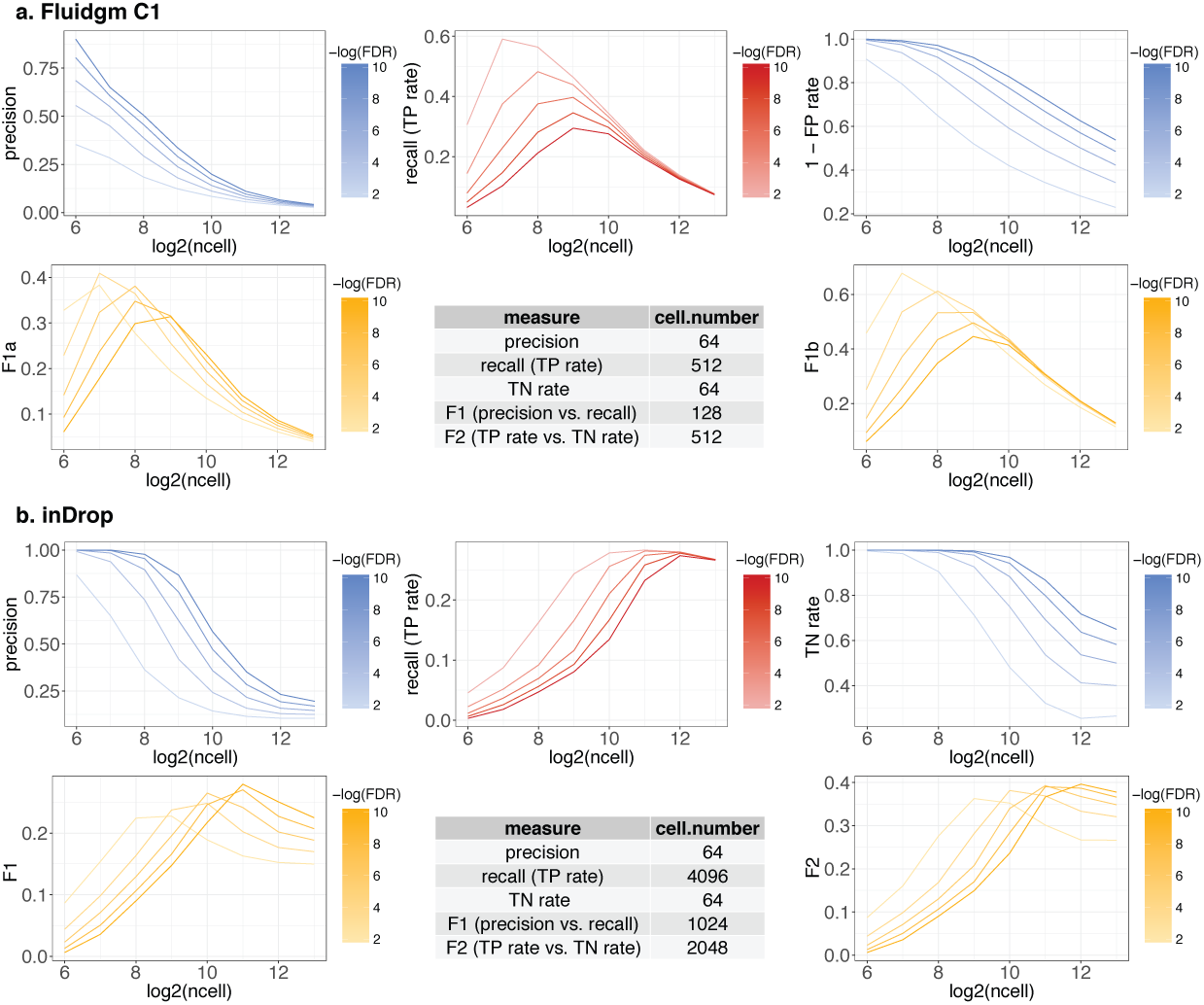
Power analysis for DE studies comparing astrocytes and oligodendrocytes (scenario 1). The thresholds on the false discovery rates (FDRs) to identify DE genes are denoted in the color legends. The table summarizes the optimal cell number according to each metric. **a:** the Fluidigm C1 protocol; **b:** the inDrop protocol.

In scenario (2), the constraint is the total sequencing depth of one experiment with at least two cell states, and the goal is to determine the optimal total cell number for that experiment given a criterion in DE analysis. scDesign simulates a new count matrix of each cell state based on a real count matrix from the same state, with pre-specified total sequencing depth, total cell number, and cell proportions of the two cell states of interest. We applied scDesign to evaluate the designs of 12 example experiments (Table S5). In every experiment, we set the sequencing depth to 100 million reads and considered six cell numbers: 512, 1024, 2048, 4096, 8192, and 16, 384. We estimated the cell proportions of the two cell states from the corresponding real data (Table S5). In practical applications of scDesign, the cell state proportions can be inferred from pilot studies, public data, or literature (Macosko et al., 2015; Gierahn et al., 2017).

In contrast to scenario (1), the optimal total cell number in scenario (2) depends on an additional factor: the cell state proportions, aside from the technical and biological factors we have discussed. The two cell states of interest may present in various proportions depending on biological conditions and experimental protocols, and larger cell state proportions in general reduce the demand of a larger total cell number. For example, the estimated cell state proportions of astrocytes and oligodendrocytes in a human brain sample are 19.2% and 14.9%, respectively (Darmanis et al., 2015), and 1024 cells are needed to maximize the recall with Fluidigm C1 (Figure S11). In a mouse visual cortex sample, however, the estimated proportions of the same two cell types are 8.8% and 13.1%, respectively, and 16, 384 cells are required to achieve the highest recall with inDrop (Figure S11). Given an experimental protocol, the optimal total cell number depends on both the two cell state proportions and the magnitude of gene expression differences between the two cell states. For example, the proportions of CD4 cells, CD8 cells, and B cells in a human peripheral blood mononuclear sample are 17.2%, 10.2%, and 7.3%, respectively (Gierahn et al., 2017). Two important facts about this experiment are: first, the proportion of CD8 cells is higher than the proportion of B cells; second, the magnitude of gene expression differences is larger between CD4 and B cells than between CD4 and CD8 cells. With the Seq-Well protocol, the DE analysis of CD4 vs. B cells only needs 4, 096 cells to achieve the highest F1 score. On the other hand, the DE analysis of CD4 vs. CD8 requires 16, 384 cells to maximize the F1 score (Figure S12). To further assess the effects of cell state proportions on DE analysis, we synthesized CD4 and B cells with multiple hypothetical cell proportions: 10%, 20%, 30%, and 40% (Figure S13), among which the mixture of 40% B cells and 20 *-* 30% CD4 cells led to the minimum cell number required to maximize the recall and precision. Determining the optimal cell state proportions given a total cell number is especially useful when the cell states of interest can be enriched by fluorescence-activated cell sorting (Jaitin et al., 2014) or flow cytometry (Yen-Rei et al., 2016) before the sequencing step.

### 3.3 scDesign demonstrates reproducibility across studies

In addition to evaluating the results of scDesign across different cell types and scRNA-seq protocols, we also analyzed the experimental designs of the same cell types and protocols but different datasets, in attempt to assess the reproducibility of scDesign.

First, we applied scDesign to optimize the pairwise DE analysis between the oligodendrocyte precursor cells (OPC) and three other brain cell types: differentiation-committed oligodendrocyte precursors (COP), myelin-forming oligodendrocytes (MFO), and newly formed oligodendrocytes (NFO). Two real datasets were collected for each cell type, and the two datasets were generated using the Fluidigm C1 protocol but from different brain regions: dorsal horn and hypothalamus (Marques et al., 2016). We applied scDesign in scenario (1), assuming a total sequencing depth of 50 million reads for each cell type. In each experiment, we assumed that the libraries of the two cell types have the same number of cells, and we considered five candidate cell numbers per cell type: 64, 128, 256, 512, and 1024. The experimental design based on the two brain regions lead to highly consistent results. Both designs show that in a DE analysis between OPCs and COPs, the optimal number of each cell type is 64 if selected by precision or true negative rate, 512 by F2 score, and 1024 by recall or F1 score (Figure 3); to better compare OPCs and MFOs or NFOs, the optimal number of each cell type is 64 if selected by precision, recall, or true negative rate, 128 by F2 score, and 64 by F1 score (Figure S14). In fact, not only do the two designs identify the same optimal cell number in each case, but they also reveal highly consistent trends about how DE accuracy changes as the number of sequenced cells increases (Figures 3, S14). This example demonstrates the reproducibility of scDesign when taking input data from biological replicates.

**Fig. 3.**
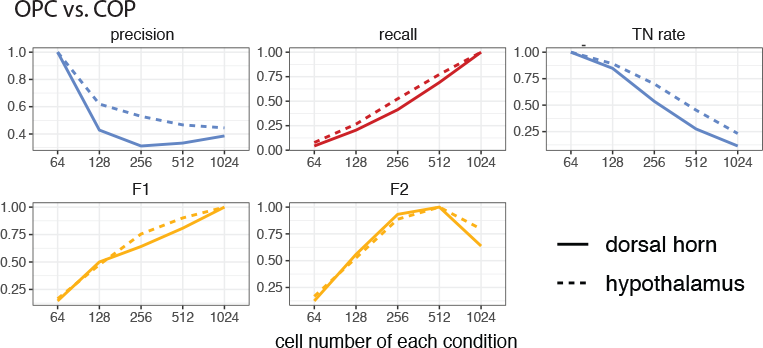
Reproducibility of scDesign based on data from different brain regions. The DE studies compare OPCs and COPs based on scRNA-seq data from two brain regions: dorsal horn and hypothalamus. When identifying the DE genes, the threshold set on the FDR rate is 10^−10^. The *y*-axis of each line are divided by the maximum value of that line for normalization.

Second, we applied scDesign to optimize the pairwise DE analysis between three retina cell types: muller glia, amacrine, and rods. Two real datasets were collected for each cell type, and the two datasets were generated using the Drop-seq protocol in two independent studies (Macosko et al., 2015; Shekhar et al., 2016). We applied scDesign in scenario (1), assuming a total sequencing depth of 100 million reads for each cell type. In each experiment, we assumed that the libraries of the two cell types have the same number of cells and considered six candidate cell numbers per cell type: 64, 128, 256, 512, 1024, and 2048. The experimental design based on the single-cell studies lead to highly similar results (Figure S15). As in the first example, both designs identified the same optimal cell number regardless of the DE criterion used. The only exception was in the comparison between the muller glia and rods: using the Macosko *et al*. data, the best cell number is 512 by F1 score and 1024 by F2 score, while using the Shekhar *et al*. data, the best cell number is 1024 by F1 score and 512 by F2 score. Such discrepancy is not suprising since we only evaluated a few candidate cell numbers, and the two input datasets inevitably differ in qualities as they came from two studies. Overall, this example demonstrates the reproducibility of scDesign when taking input data from independent studies to design scRNA-seq experiments.

### 3.4 scDesign assists scRNA-seq method development

In addition to assisting single-cell experimental design, scDesign can also simulate scRNA-seq data to benchmark various computational methods for differential gene expression analysis, single cell clustering analysis, gene expression dimension reduction, etc. Due to excess zeros resulting from dropout events and the fact that each gene’s expression level in each cell is only measured once, the ground truth of individual genes’ expression levels in single cells cannot be accurately estimated from scRNA-seq data. Also, cellular identities of individual cells are difficult to pre-determine in most experiments. Lacking the aforementioned ground truth encumbers the development of computational methods to decipher information from scRNA-seq data. Direct evaluation of computational methods relies on experimental validation, which is often unavailable for computationalists, and indirect interpretation from downstream analysis is used instead as a not-so-ideal substitute. Empowered by its ability to generate synthetic scRNA-seq data that well mimic real scRNA-seq data and have ground truth information, scDesign provides a flexible framework to benchmark computational methods for various scRNA-seq data analysis tasks.

We first demonstrated the application of scDesign to evaluating DE methods. We considered a baseline DE method, i.e., the two-sample *t* test, and four DE methods (MAST (Finak et al., 2015), SCDE (Kharchenko et al., 2014), DESeq2 (Love et al., 2014), and edgeR (Robinson et al., 2010)) specifically designed for scRNA-seq data. Here both DESeq2 and edgeR denote their single-cell-adapted versions, where gene expression values are weighted by the weights estimated from a zero inflated negative Binomial model before the statistical testing step (Van den Berge et al., 2017). We evaluated scDesign using real scRNA-seq data of six cell types: dendrocytes (Smart-seq2, 63.6% zero count), oligodendrocytes (Fluidigm C1, 62.9% zero count), interneurons (inDrop, 75.3% zero counts), retinal ganglions (Drop-Seq, 78.3% zero counts), enterocytes (10× Genomics, 82.0% zero counts), and natural killer cells (Seq-Well, 88.0% zero counts) (Table S1). Based on the real data of each cell type, we simulated a pair of count matrices, with one matrix representing the given cell type and the other including up-regulated and down-regulated genes (Supplementary Information). In the first setting, we set the percentage of DE genes to 5% and sampled the fold changes of those DE genes’ expression values uniformly from the interval [2, 5] (Supplementary information). Then we evaluated the performance of the five DE methods by comparing the areas under their precision-recall curves (Figure S16). With Smart-seq2 and Fluidigm C1, MAST and SCDE were the only two methods that achieved better accuracy than the two-sample *t* test, but overall the three methods had comparable precision and recall. With inDrop and 10× Genomics, edgeR became the best DE method, followed by MAST and SCDE. With Drop-seq and Seq-Well, the most accurate method was SCDE, and the baseline two-sample *t* test had poor performance. These simulation results suggest that scRNA-seq data from the 10×, inDrop, Drop-seq, and Seq-Well protocols need more specialized statistical modeling in the DE analysis, compared with Smart-seq2 and Fluidigm C1. In the second setting, we set the percentage of up-regulated and down-regulated genes in each comparison to 10% and sampled the fold changes of these DE genes uniformly from the interval [4, 5]. Since the magnitude of fold changes increased, the DE methods overall demonstrated improved accuracy (Figure S17), but the relative accuracy of the five DE methods was consistent with that under the first setting.

We next demonstrated the application of scDesign to comparing dimension reduction methods. We considered four methods: principal component analysis (PCA), *t*-distributed stochastic neighbor embedding (tSNE) (Maaten and Hinton, 2008), independent component analysis (ICA) (Hyvärinen and Oja, 2000), and ZINB-WaVE (Risso et al., 2018). We evaluated scDesign based on the same real scRNA-seq data used in the comparison of DE methods. Based on the real data of each cell type, we simulated a set of synthetic count matrices, representing multiple cell states following a differentiation path (Supplementary Information). For the Smart-seq2 and Fluidigm C1 protocols, we simulated four cell states with two states each having 80 cells and the other two each having 50 cells. For the other four protocols, we simulated five cell states with two states each having 300 cells and the other three each having 100 cells. In each case, we first simulated the cell state at the starting point of differentiation based on the real data, and then we simulated each of the three subsequent cell states with 1% of up-regulated and down-regulated genes from its previous state. In addition, we sampled the fold changes of those DE genes’ expression values uniformly from [2, 5]. Among the four dimension reduction methods, ZINB-WaVE had the best performance in grouping cells into biologically meaningful clusters based on the C1 data, followed by the tSNE method (Figure S18). Based on the Smart-Seq2 data, PCA had the best performance in the 2D space, followed by ZINB-WaVE (Figure S19). However, the comparison results were different for droplet-based protocols. The tSNE method led to the most accurate cell clusters for the Drop-seq, inDrop, 10×, and Seq-Well protocols, followed by ICA or PCA (Figures S20-S23). In spite of the clustering performance, another factor worth noting is that PCA, ICA, and ZINB-WaVE generate comparable cell-cell distances after dimension reduction, but tSNE does not. The above results demonstrate the capacity of scDesign in helping developers evaluate competing computational methods for the same purpose (e.g., DE analysis or dimension reduction), and in assisting users to select the appropriate method for analyzing scRNA-seq data from a specific protocol.

## 4 Discussion

The scRNA-seq technologies have become an essential tool for studying various biological and biomedical problems, but one unresolved challenge is how to balance the trade-off between exploring the depth or breadth of transcriptome information in experimental design. We introduce scDesign, the first statistical and computational simulator that enables rational and practical scRNA-seq experimental design. By integrating statistical assumptions and real scRNA-seq datasets from public repositories into its generative framework, scDesign is able to mimic the real experimental processes and simulate synthetic scRNA-seq datasets that well capture gene expression characteristics in real data. In addition, scDesign is a flexible and reproducible simulator that is capable of modeling protocol-specific scRNA-seq data generated under multiple biological and experimental conditions. We conducted a comprehensive comparison of scDesign and four other scRNA-seq simulation methods based on datasets from 17 different cell types and six experimental protocols. The comparison suggests that scDesign generates synthetic data with the largest resemblance to real scRNA-seq data regardless of cell types and protocols.

Using its simulated data, scDesign performs power analysis on differential gene expression analysis to provide a quantitative and objective standard for designing future experiments. In the context of differential gene expression analysis between two cell states, scDesign suggests an optimal cell number given a fixed sequencing depth, in the trade-off between a deeper sequencing of a smaller number of cells or a shallower sequencing of a larger number of cells. Specifically, we demonstrated the application of scDesign in two scenarios, where cells from the two states are sequenced as two separate libraries or as one pooled library. We evaluated the experimental designs for 14 and 12 scRNA-seq studies under the two scenarios, respectively. Our results for the first time demonstrate how the optimal experimental design for DE analysis depends on the scRNA-seq protocol and the intra and inter cell state transcriptome heterogeneity. In addition, our results revealed a general phenomenon that a deeper sequencing of a smaller number of cells leads to a higher precision in DE analysis. In contrast to the precision, maximizing the recall of DE analysis requires finding a balance between the cell-wise sequencing depth and the cell number, because our results show that the recall first increases and then decreases as we increase the cell number with the total sequencing depth fixed. scDesign enables researchers to design effective scRNA-seq experiments without pre-experimental costs in an objective manner, for example, guided by the expected power in downstream DE analysis. In addition, we demonstrate that scDesign leads to reproducible experimental design for target cell states given data generated in different studies.

Aside from enhancing future experimental design, another main contribution of scDesign is to assist computational method development for scRNA-seq. Since large-scale benchmark data are not yet available in the field, computationalists typically rely on scRNA-seq datasets from public repositories to test and evaluate new methods and algorithms. However, quality control and normalization of real data are themselves ongoing research questions, making the evaluation results in many method papers not comparable nor reproducible (McCarthy et al., 2017; Ziegenhain et al., 2017). To tackle this challenge, scDesign allows users to generate synthetic scRNA-seq datasets with user-specified experimental protocols, sequencing depths, cell states, cell numbers, as well as pre-specified differentially expressed genes. Given that scDesign generates synthetic data with known truth and well mimicking real data, users can leverage its synthetic data to comprehensively evaluate computational and statistical methods in a flexible, reproducible, and comparable way. For example, we compared five DE methods (the two-sample *t* test, MAST, SCDE, DESeq2, and edgeR) and four dimension reduction methods (PCA, tSNE, ICA, and ZINB-WaVE) using synthetic data generated by scDesign. Those comparison results provide useful guidance for researchers to select the most appropriate computational method to analyze real data.

We expect scDesign to assist scRNA-seq experimental design for a vast array of available experimental protocols. scDesign incorporates real scRNA-seq data into its statistical framework to make flexible decisions based on the protocol and cell states used in the target study. If the real data of the two cell states are not generated from the same experiment, it is recommended to first correct the batch effect before applying scDesign (Haghverdi et al., 2018; Butler et al., 2018). To extend scDesign’s ability to evaluate experimental designs for cell states whose scRNA-seq data are not yet publicly available, a future direction is to incorporate bulk RNA-seq data of the same type as a surrogate to estimate the gene expression parameters. Otherwise, pilot experiments need to be conducted to collect data for experimental design, which is also a widely adopted practice (Chatterjee et al., 2018). Another future extension of scDesign is to find the optimal design in the context of other types of downstream analyses besides the differential gene expression analysis, such as the detection of novel cell sub-types or the recovery of temporal transcriptome trajectories (Dumitrascu et al., 2018a). For instance, we may jointly learn the proportions and the gene expression profiles of multiple cell states from real scRNA-seq data and use them as input into our simulation framework to evaluate how the power of detecting rare cell types changes with experimental parameters. Given time-series scRNA-seq data, the scDesign framework can be modified to conduct ANOVA or more advanced statistical analysis to objectively select cell numbers for multiple time points. It is also possible to generalize the simulation framework of scDesign to account for more complex trajectories in the cell differentiation process (Papadopoulos et al., 2019; Cannoodt et al., 2019). We expect scDesign to be an effective bioinformatic tool that assists rational scRNA-seq experiment design based on specific research goals and benchmarks competing scRNA-seq computational methods.

