## Supplementary materials for "A statistical simulator scDesign for rational scRNA-seq experimental design"

##### Simulating multiple count matrices following a differentiation path

Given a real dataset with $I$ genes and $J_0$ cells, the goal of this section is to generate $G$ $(G \geq 2)$ new count matrices, each of which has $I$ genes, $J$ synthetic cells, and a total of $S$ reads. The synthetic data should represent $G$ cell states following a specified differentiation path with known DE genes, such that these data serve as a good basis for benchmarking single-cell data analysis and method development. When generating the $G$ synthetic count matrices, we assume that the $G$ cell states follow a differentiation path, with a $p_{\text{up}}$ proportion of up-regulated genes and a $p_{\text{down}}$ proportion of down-regulated genes from state $g$ to state $g+1$ $(g = 1, \ldots, G-1)$.

1. Estimate parameters from real scRNA-seq data.

As described in **Simulating a single count matrix**, from the real count matrix $\boldsymbol{X}^{\text{real}}_{I \times J_0}$, we obtained the following parameter estimates: (1) the mean $\hat{\mu}_s$ and the standard deviation $\hat{\sigma}_s$ of the Normal distribution used to model the cell library sizes; (2) the cell-wise dropout rates $\hat{q}_{01}, \ldots, \hat{q}_{0J_0}$; (3) the gene-wise dropout rate $\hat{\lambda}_{0i}$, mean $\hat{\mu}_{0i}$, and standard deviation $\hat{\sigma}_{0i}$ of gene $i$, $i = 1, \ldots, I$. A Gamma distribution was used to fit the estimated gene mean expression $\hat{\mu}_{01}, \ldots, \hat{\mu}_{0I}$ and the estimated shape and scale parameters are denoted as $\hat{k}_0$ and $\hat{\theta}_0$. The above parameter estimates were used to simulate the expression parameters of state $1$, while the parameters of state $g+1$ depended on the parameters of its previous state $g$.

2. Simulate gene mean expression values of the $G$ states.

In this step, we simulated the log-scale mean gene expression values under each cell state, without considering dropout events. We assumed that from state $g$ to state $g+1$, the proportions of up-regulated and down-regulated genes were $p_{\text{up}}$ and $p_{\text{down}}$, respectively. The fold changes of gene mean expression levels were independently and uniformly distributed within $[f_l, f_u]$.

We used $\mu_i^g$ to denote the mean expression of gene $i$ in cell state $g$. For cell state $1$, we simulated $\mu_i^1$ from the Gamma distribution: $\mu_i^1 \overset{\text{i.i.d.}}{\sim} \text{Gamma}(\hat{k}_0, \hat{\theta}_0)$, $i = 1, \ldots, I$. Then given $\mu_1^g, \ldots, \mu_I^g$, we simulated $\mu_1^{g+1}, \ldots, \mu_I^{g+1}$ $(g = 1, \ldots, G-1)$ as follows.

(a) We simulated the number of up-regulated genes $n^g_{\text{up}}$, and the number of down-regulated genes $n^g_{\text{down}}$ from a Multinomial distribution:

$$\left(n^g_{\text{up}}, n^g_{\text{down}}, I - n^g_{\text{up}} - n^g_{\text{down}}\right) \sim \text{Multinomial}\left(I, (p_{\text{up}}, p_{\text{down}}, 1 - p_{\text{up}} - p_{\text{down}})\right) .$$

(b) We randomly drew the $n^g_{\text{up}} + n^g_{\text{down}}$ DE genes from the gene population $\{1, \ldots, I\}$ without replacement and denoted

$$d_i^g = \begin{cases} 1, & \text{if gene } i \text{ is up-regulated} \\ -1, & \text{if gene } i \text{ is down-regulated} \\ 0, & \text{otherwise} \end{cases} .$$

(c) We simulated $\mu_I^{g+1}$, the mean expression of gene $i$ in state $g+1$, as follows:

$$\mu_i^{g+1} = \begin{cases} \mu_i^g + \log_{10} f_i^g & \text{if } d_i^g = 1 \\ \mu_i^g - \log_{10} f_i^g & \text{if } d_i^g = -1 \\ \mu_i^g, & \text{otherwise} \end{cases} ,$$

where $f_i^g \overset{\text{i.i.d}}{\sim} \text{Uniform}[f_l, f_u])$.

3. Simulate the count matrices.

With the mean gene expression $\mu_1^g, \ldots, \mu_I^g$, we simulated the count matrix $\boldsymbol{X}^{\text{syn},g}$ under each state $g$ independently following steps 2-4 in **Simulating a single count matrix**. Please note that we estimated and simulated other cell-wise and gene-wise parameters also by following **Simulating a single count matrix**. We kept the estimated parameters the same for all the cell states, and we simulated the cell-wise parameters of synthetic cells independently across all the states.

#### Comparison of different simulation methods

The splat, Lun, and scDD simulation methods were implemented using the R package `splatter` version 1.3.3.9010. The powsimR method was implemented using the R package `powsimR` version 1.1.0. The scDesign method was implemented using the R package `scDesign` version 1.0.0.

We denote a $\log 10$-transformed count matrix as $\boldsymbol{X}_{I \times J}$, with rows representing genes and columns representing cells. For gene $i$ ($i = 1, \ldots, I$), we define its count mean $T_i^1 = \frac{1}{J}\sum_{j=1}^J X_{ij}$, count variance $T_i^2 = \frac{1}{J-1}\sum_{j=1}^J (X_{ij} - T_i^1)^2$ , coefficient of variance (cv) $T_i^3 = \frac{\sqrt{T_i^2}}{T_i^1}$, and gene-wise zero proportion $T_i^4 = \frac{1}{J}\sum_{j=1}^J \mathbb{I}\{X_{ij} = 0\}$. For each cell $j$ ($j = 1, \ldots, J$), we calculated its library size $T_j^5 = \sum_{i=1}^I X_{ij}$ and cell-wise zero proportion $T_j^6 = \frac{1}{I}\sum_{i=1}^I \mathbb{I}\{X_{ij} = 0\}$. For each real $\log 10$-transformed matrix, we calculated the values of the six statistics

$$\left\{ \left( T_i^{1,\text{real}}, T_i^{2,\text{real}}, T_i^{3,\text{real}}, T_i^{4,\text{real}}, T_j^{5,\text{real}}, T_j^{6,\text{real}} \right) : i = 1, \ldots, I, j = 1, \ldots, J \right\}$$

and denote the resulting empirical distribution of the $k$-th statistic as $F^k$, $k = 1, , \ldots, 6$. For each synthetic $\log 10$-transformed matrix, we also calculated the values of the six statistics

$$\left\{ \left( T_i^{1,\text{syn}}, T_i^{2,\text{syn}}, T_i^{3,\text{syn}}, T_i^{4,\text{syn}}, T_j^{5,\text{syn}}, T_j^{6,\text{syn}} \right) : i = 1, \ldots, I, j = 1, \ldots, J \right\}$$

and denote the resulting empirical distribution of the $k$-th statistic as $G^k$, $k = 1, , \ldots, 6$. Finally, to evaluate the quality of the synthetic data, we calculated the Kolmogorov-Smirnov (KS) distance between $F^k$ and $G^k$ is calculated as

$$D(F^k, G^k) = \sup_x |F^k(x) - G^k(x)|,\ x \in \mathbb{R}\,.$$

**Table S1:** Details of real scRNA-seq data. Cell type information is from the original work that published the corresponding data.

| data | accession | species | cell type | protocol |
|---|---|---|---|---|
| 1 | GSE94820 | human | dendrocyte1 (165)<br>dendrocyte2 (94)<br>monocyte2 (163) | Smart-Seq2 |
| 2 | GSM1626793 | mouse | bipolar (919)<br>cones (241)<br>rods (3746)<br>retinal ganglion (70) | Drop-seq |
| 3 | GSE92332 | mouse | goblet (510)<br>stem (1267)<br>tuft (166) | 10x |
| 4 | GSE67835 | human | astrocyte (49)<br>neuron (122)<br>oligodendrocyte (38) | Fluidigm C1 |
| 5 | GSE102827 | mouse | excitatory (1040)<br>neuron (116)<br>oligodendrocyte (286)<br>astrocyte (189) | inDrop |
| 6 | GSM2486333 | human | natural killer (471)<br>CD4 (634)<br>CD8 (269)<br>B cell (376) | Seq-Well |

**Table S2:** The five metrics used to evaluate accuracy of DE analysis.

| metric | abbreviation | meaning |
|---|---|---|
| recall (true positive rate) | recall (TP) | ability of a method to identify all true DE genes |
| precision | precision | ability of a method to identify only true DE genes |
| 1-false positive rate | 1-FP | |
| harmonic mean of recall and precision | F1 | combined ability of a method to identify all and only true DE genes |
| harmonic mean of TP and 1-FP | F2 | |

**Table S3:** The optimal cell numbers in experimental design (scenario 1) using t test. Five DE accuracy measures are calculated in every comparison: precision, recall, TN (true negative rate), F1, and F2. For each measure, the smallest cell number that leads to the highest accuracy is recorded as the optimal number.

| protocol | cell type 1 | cell type 2 | precision | recall | TN | F1 | F2 |
|---|---|---|---|---|---|---|---|
| Smart-Seq2 | dendrocyte1 | monocyte1 | 64 | 256 | 64 | 128 | 128 |
| Smart-Seq2 | dendrocyte1 | dendrocyte2 | 64 | 512 | 64 | 256 | 512 |
| Drop-seq | cone | retinal ganglion | 64 | 1024 | 64 | 512 | 512 |
| Drop-seq | cone | rod | 64 | 2048 | 64 | 1024 | 512 |
| 10x | tuft | goblet | 64 | 2048 | 64 | 1024 | 4096 |
| 10x | tuft | stem | 64 | 4096 | 64 | 2048 | 4096 |
| C1 | neuron | astrocyte | 64 | 512 | 64 | 128 | 512 |
| C1 | neuron | oligodendrocyte | 64 | 512 | 64 | 128 | 512 |
| C1 | astrocyte | oligodendrocyte | 64 | 512 | 64 | 128 | 512 |
| inDrop | astrocyte | oligodendrocyte | 64 | 4096 | 64 | 1024 | 2048 |
| inDrop | excitatory | interneuron | 64 | 4096 | 64 | 2048 | 4096 |
| inDrop | excitatory | oligodendrocyte | 64 | 1024 | 64 | 128 | 512 |
| Seq-Well | CD4 | B cell | 64 | 2048 | 64 | 512 | 512 |
| Seq-Well | CD4 | CD8 | 64 | 8192 | 64 | 8192 | 8192 |

**Table S4:** The optimal cell numbers in experimental design (scenario 1) using MAST. Five DE accuracy measures are calculated in every comparison: precision, recall, TN (true negative rate), F1, and F2. For each measure, the smallest cell number that leads to the highest accuracy is recorded as the optimal number.

| protocol | cell type 1 | cell type 2 | precision | recall | TN | F1 | F2 |
|---|---|---|---|---|---|---|---|
| Smart-Seq2 | dendrocyte1 | monocyte1 | 64 | 128 | 64 | 64 | 128 |
| Smart-Seq2 | dendrocyte1 | dendrocyte2 | 64 | 256 | 64 | 128 | 256 |
| Drop-seq | cone | retinal ganglion | 64 | 1024 | 64 | 512 | 1024 |
| Drop-seq | cone | rod | 64 | 4096 | 64 | 4096 | 4096 |
| 10x | tuft | goblet | 64 | 1024 | 64 | 512 | 1024 |
| 10x | tuft | stem | 64 | 2048 | 64 | 512 | 2048 |
| C1 | neuron | astrocyte | 64 | 512 | 64 | 128 | 512 |
| C1 | neuron | oligodendrocyte | 64 | 512 | 64 | 256 | 512 |
| C1 | astrocyte | oligodendrocyte | 64 | 256 | 8192 | 256 | 256 |
| inDrop | astrocyte | oligodendrocyte | 64 | 2048 | 64 | 1024 | 2048 |
| inDrop | excitatory | interneuron | 64 | 1024 | 64 | 256 | 1024 |
| inDrop | excitatory | oligodendrocyte | 64 | 2048 | 64 | 1024 | 2048 |
| Seq-Well | CD4 | B cell | 64 | 2048 | 64 | 1024 | 2048 |
| Seq-Well | CD4 | CD8 | 64 | 4096 | 64 | 4096 | 4096 |

**Table S5:** The optimal cell numbers in experimental design (scenario 2) using t tests. Five DE accuracy measures are calculated in every comparison: precision, recall, TN (true negative rate), F1, and F2. For each measure, the smallest cell number that leads to the highest accuracy is recorded as the optimal number. The proportions of each cell type in the cell population are listed under the cell labels.

| protocol | cell type 1 | cell type 2 | precision | recall | TN | F1 | F2 |
|---|---|---|---|---|---|---|---|
| Smart-Seq2 | dendrocyte1 16.1% | monocyte1 15.8% | 512 | 1024 | 512 | 512 | 1024 |
| Smart-Seq2 | dendrocyte1 16.1% | dendrocyte2 9.1% | 512 | 2048 | 512 | 1024 | 2048 |
| Drop-seq | cone 0.42% | retinal ganglion 0.1% | 8192 | 16384 | 16384 | 16384 | 16384 |
| Drop-seq | cone 0.42% | rod 65.6% | 2048 | 16384 | 2048 | 16384 | 16384 |
| 10x | tuft 2.3% | goblet 7.1% | 512 | 16384 | 512 | 16384 | 16384 |
| 10x | tuft 2.3% | Stem 17.6% | 1024 | 16384 | 1024 | 16384 | 16384 |
| C1 | neuron 47.8% | astrocyte 19.2% | 512 | 512 | 512 | 512 | 512 |
| C1 | astrocyte 19.2% | oligodendrocyte 14.9% | 512 | 1024 | 512 | 1024 | 1024 |
| inDrop | astrocyte 8.8% | oligodendrocyte 13.1% | 512 | 16384 | 512 | 8192 | 16384 |
| inDrop | excitatory 47.8% | interneuron 5.3% | 512 | 16384 | 512 | 8192 | 16384 |
| Seq-Well | CD4 17.2% | B cell 7.3% | 512 | 16384 | 512 | 4096 | 8192 |
| Seq-Well | CD4 17.2% | CD8 10.2% | 512 | 16384 | 512 | 16384 | 16384 |

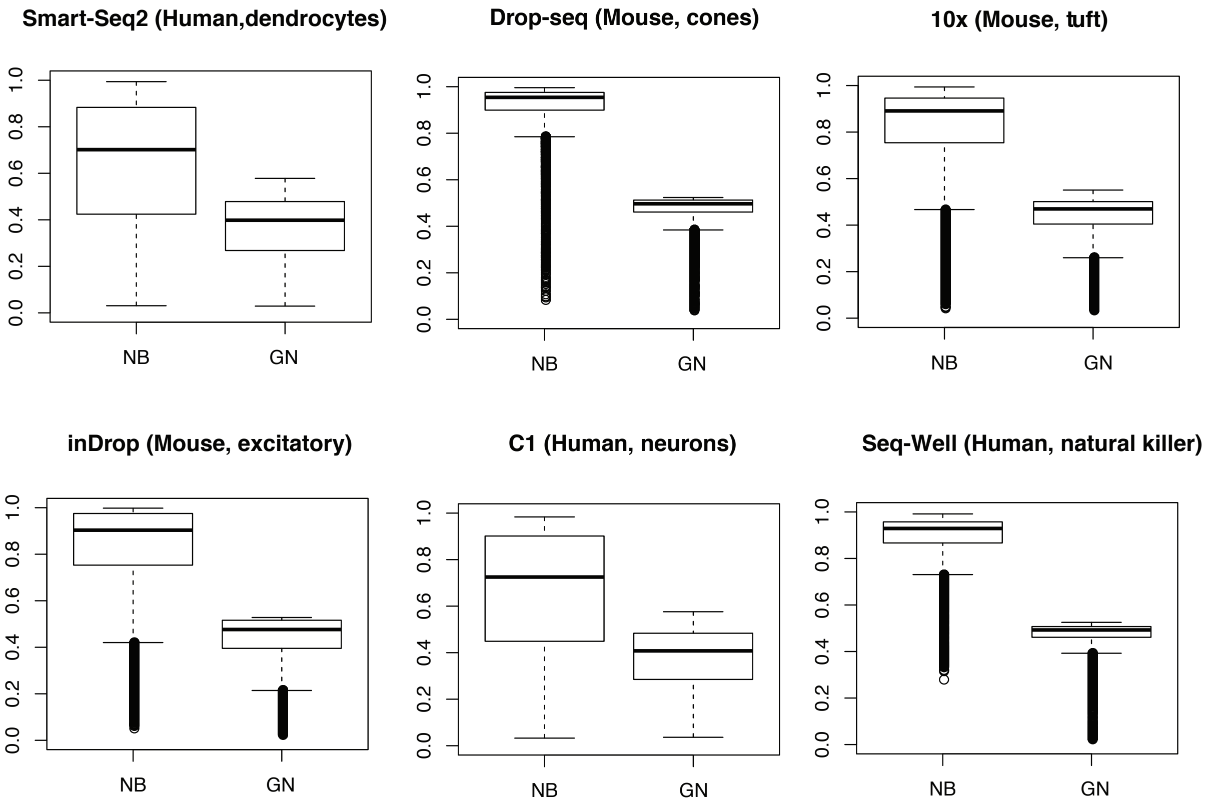


**Figure S1:** Comparison of the Negative Binomial (NB) model and the Gamma-Normal (GB) model used in scImpute. Both models are used to fit six scRNA-seq datasets from different protocols, as listed in Table S1. For each gene, the Kolmogorov-Smirnov (KS) distance between the empirical and fitted gene expression distribution is calculated and summarized in the boxplots. In all the scenarios, the GB model leads to smaller KS distances.

#### Dendrocytes subtype 1

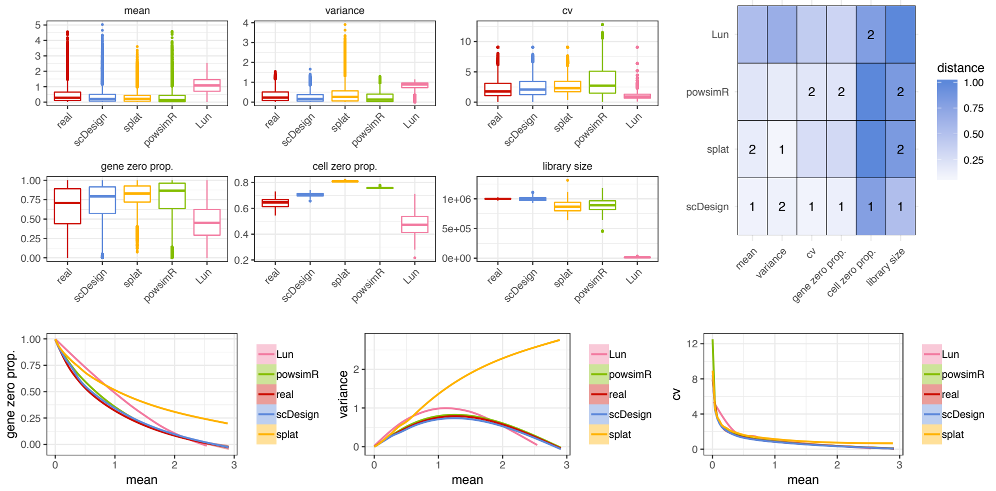


#### Dendrocytes subtype 2

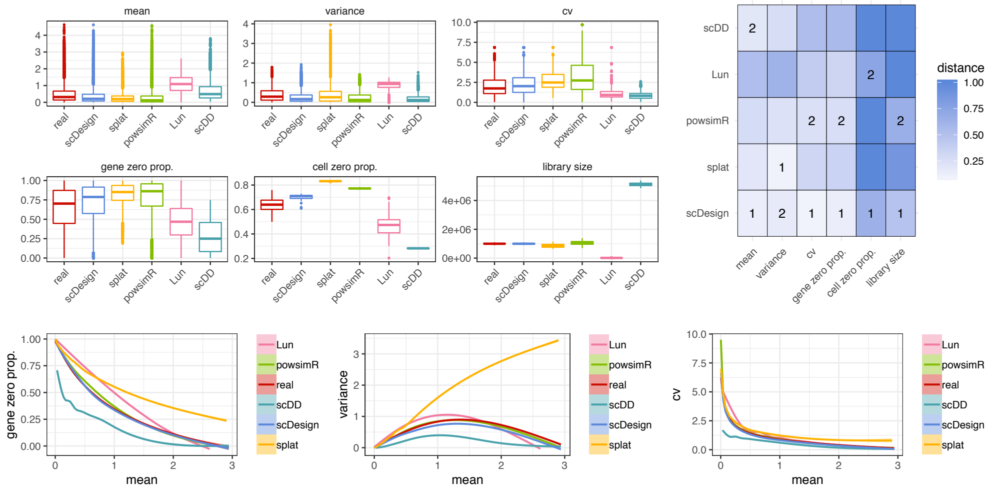


**Figure S2:** Comparison of scDesign and the other four simulation methods based on the Smart-seq2 protocol. The boxplots display the gene-wise expression mean, expression variance, expression coefficient of variation, zero proportion, and the cell-wise zero proportion and library size in both real and simulated datasets. The heatmaps display the KS distances between the six statistics in the real data and in the simulated data. The best and second best simulation methods with respect to each statistic are respectively marked with $1$ and $2$ in the heatmaps. The line plots demonstrate the empirical relationships between the key statistics in real and simulated data. Note that scDD failed to simulate data for the dendrocytes subtype1 dataset.

### Astrocytes

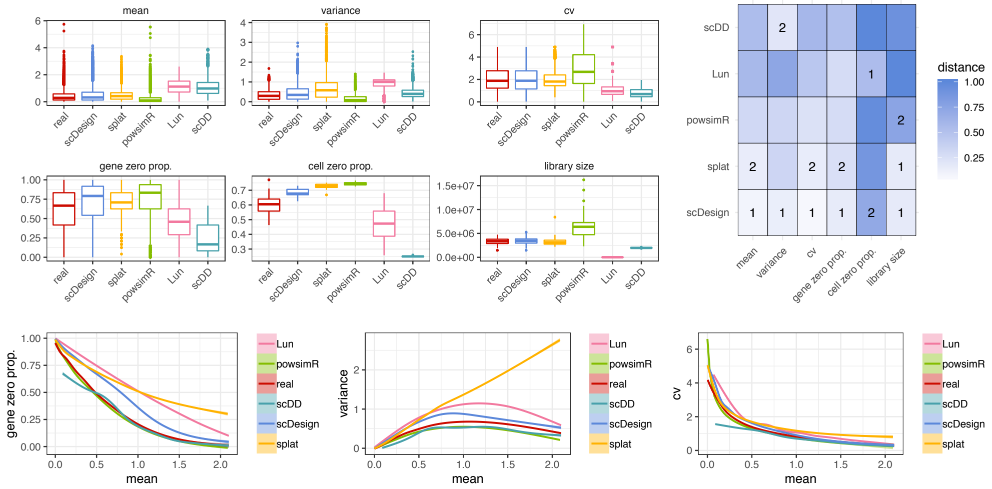


### Neuron cells

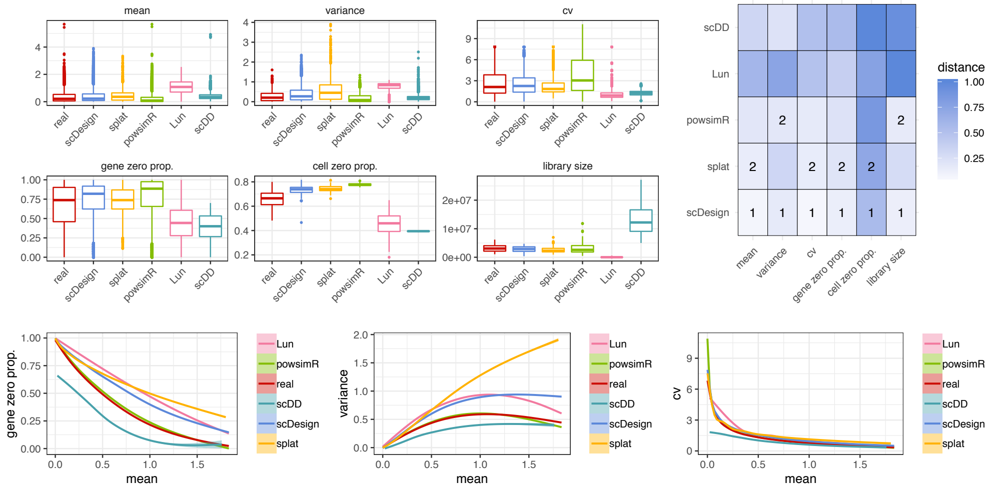


### Oligodendrocytes

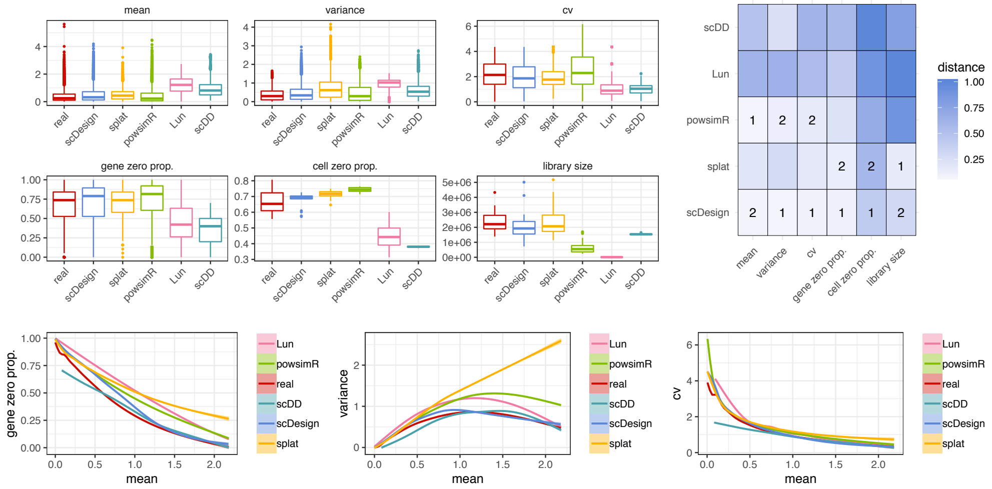


**Figure S3:** Comparison of scDesign and the other four simulation methods based on the Fluidigm C1 protocol. The boxplots display the gene-wise expression mean, expression variance, expression coefficient of variation, zero proportion, and the cell-wise zero proportion and library size in both real and simulated datasets. The heatmaps display the KS distances between the six statistics in the real data and in the simulated data. The line plots demonstrate the empirical relationships between the key statistics in real and simulated data.

### CD4 cells

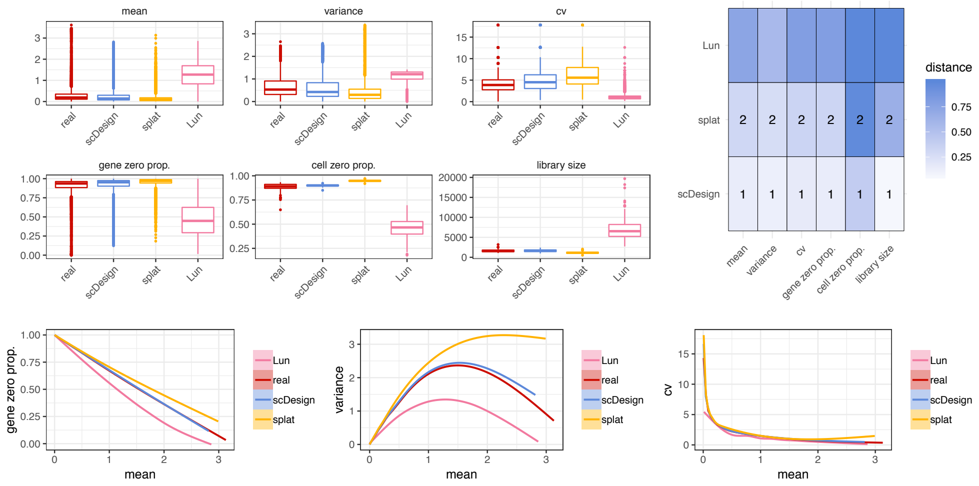


### CD8 cells

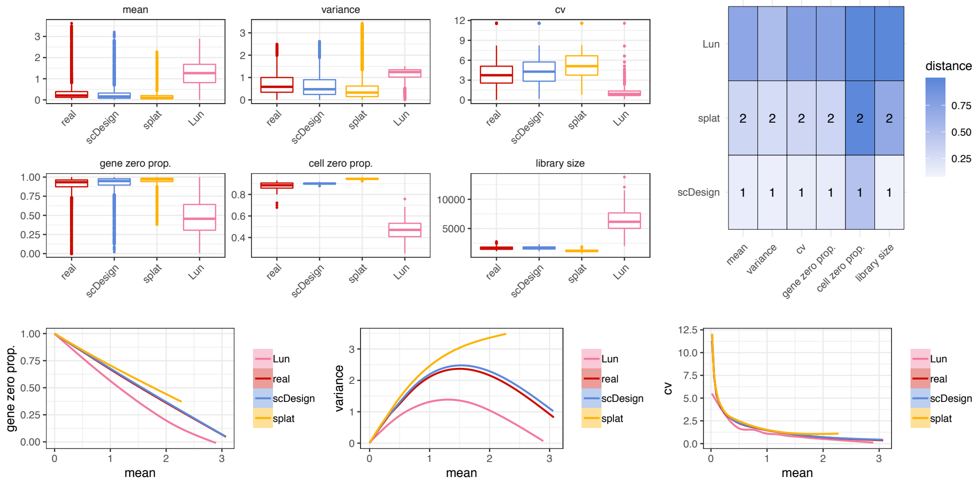


### Natural killer cells

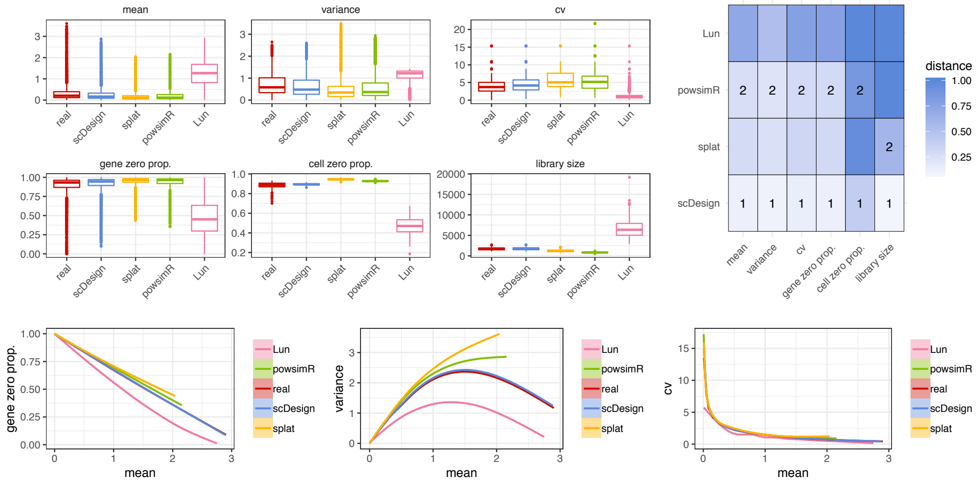


**Figure S4:** Comparison of scDesign and the other four simulation methods based on the Seqwell protocol. The boxplots display the gene-wise expression mean, expression variance, expression coefficient of variation, zero proportion, and the cell-wise zero proportion and library size in both real and simulated datasets. The heatmaps display the KS distances between the six statistics in the real data and in the simulated data. The line plots demonstrate the empirical relationships between the key statistics in real and simulated data. Note that scDD failed to simulate data for the three datasets, and powsimR failed to simulate data for the CD4 and CD8 dataset.

Goblets

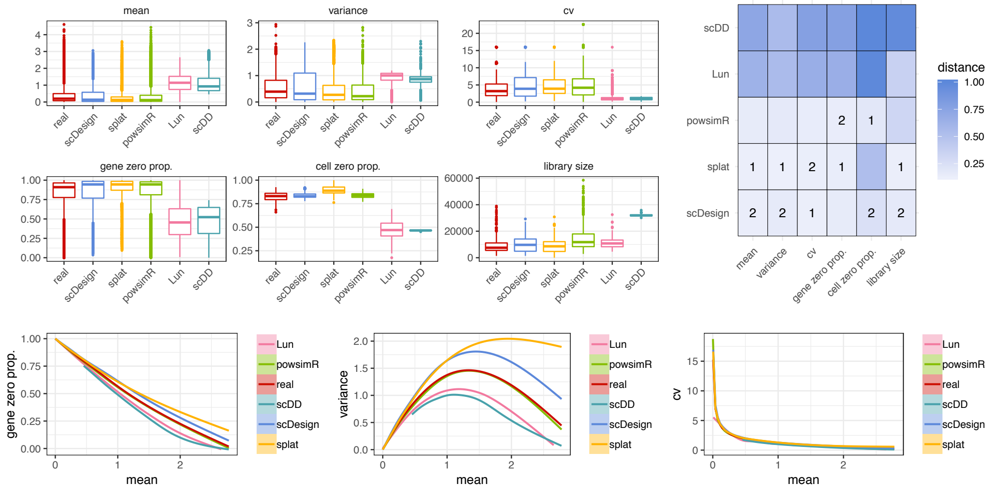


Stem cells

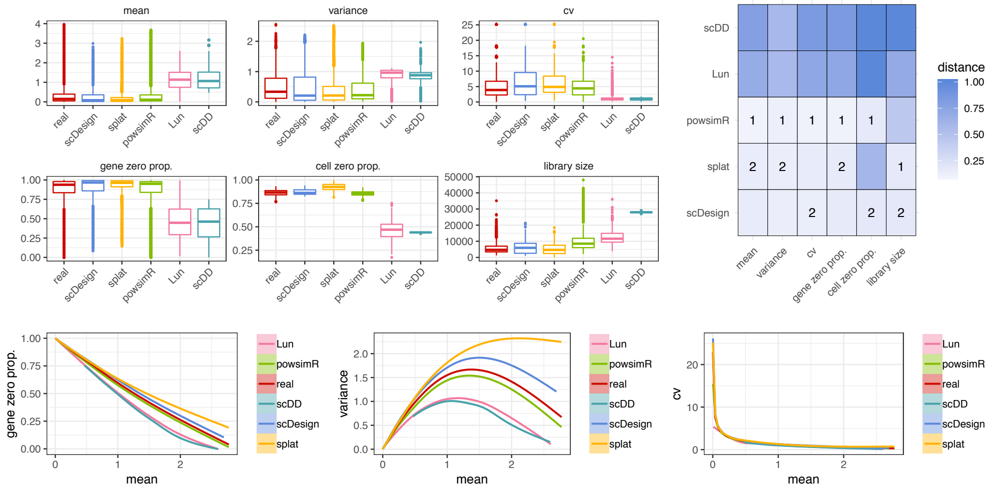


Tufts

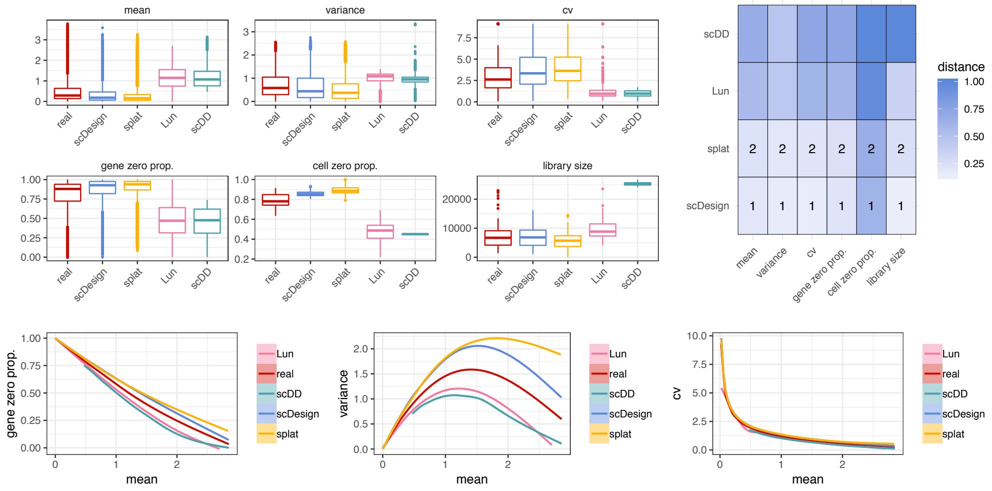


**Figure S5:** Comparison of scDesign and the other four simulation methods based on the 10x Genomics protocol. The boxplots display the gene-wise expression mean, expression variance, expression coefficient of variation, zero proportion, and the cell-wise zero proportion and library size in both real and simulated datasets. The heatmaps display the KS distances between the six statistics in the real data and in the simulated data. The line plots demonstrate the empirical relationships between the key statistics in real and simulated data. Note that powsimR failed to simulate data for the Tufts dataset.

### Bipolar cells

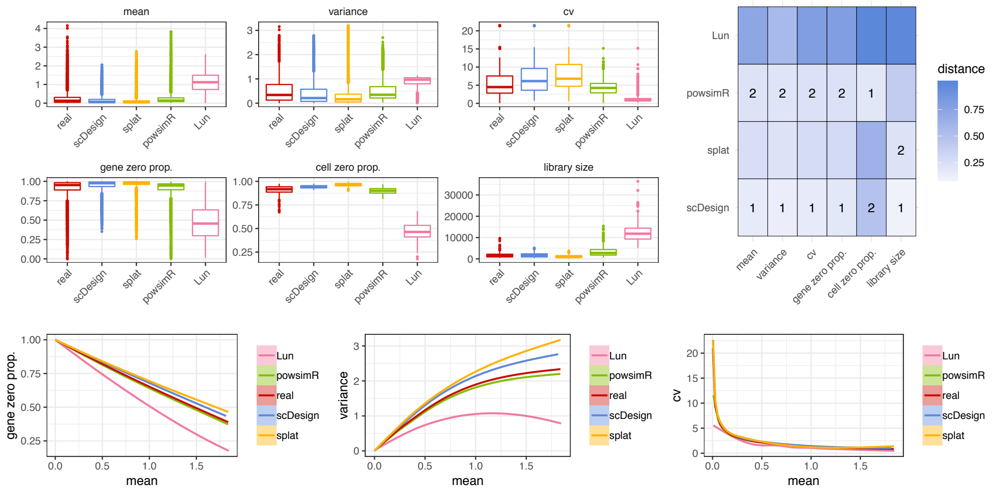


### Cones

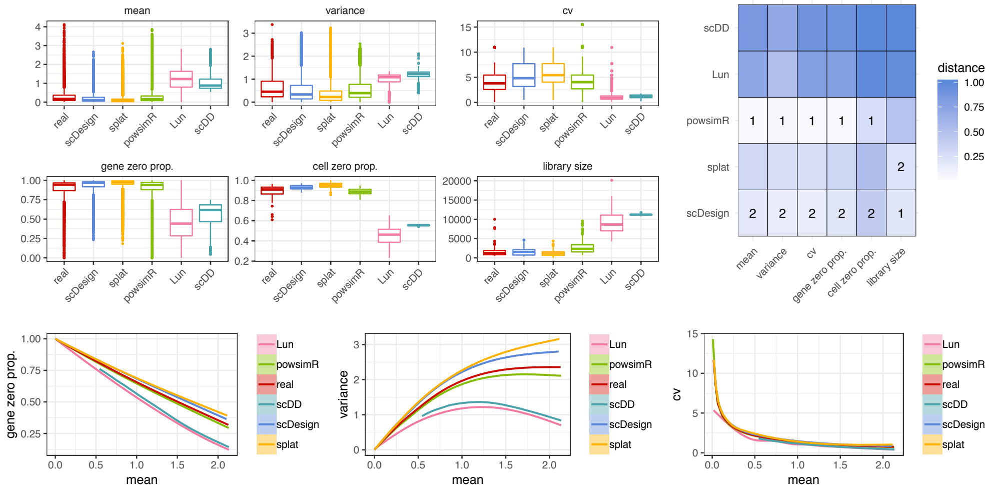


### Rods

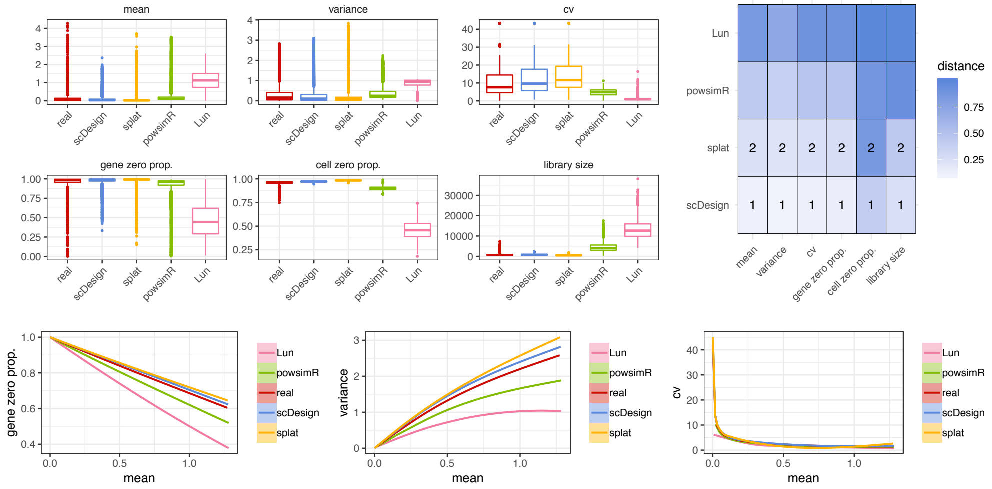


**Figure S6:** Comparison of scDesign and the other four simulation methods based on the Dropseq protocol. The boxplots display the gene-wise expression mean, expression variance, expression coefficient of variation, zero proportion, and the cell-wise zero proportion and library size in both real and simulated datasets. The heatmaps display the KS distances between the six statistics in the real data and in the simulated data. The line plots demonstrate the empirical relationships between the key statistics in real and simulated data. Note that scDD failed to simulate data for the bipolar and rod datasets.

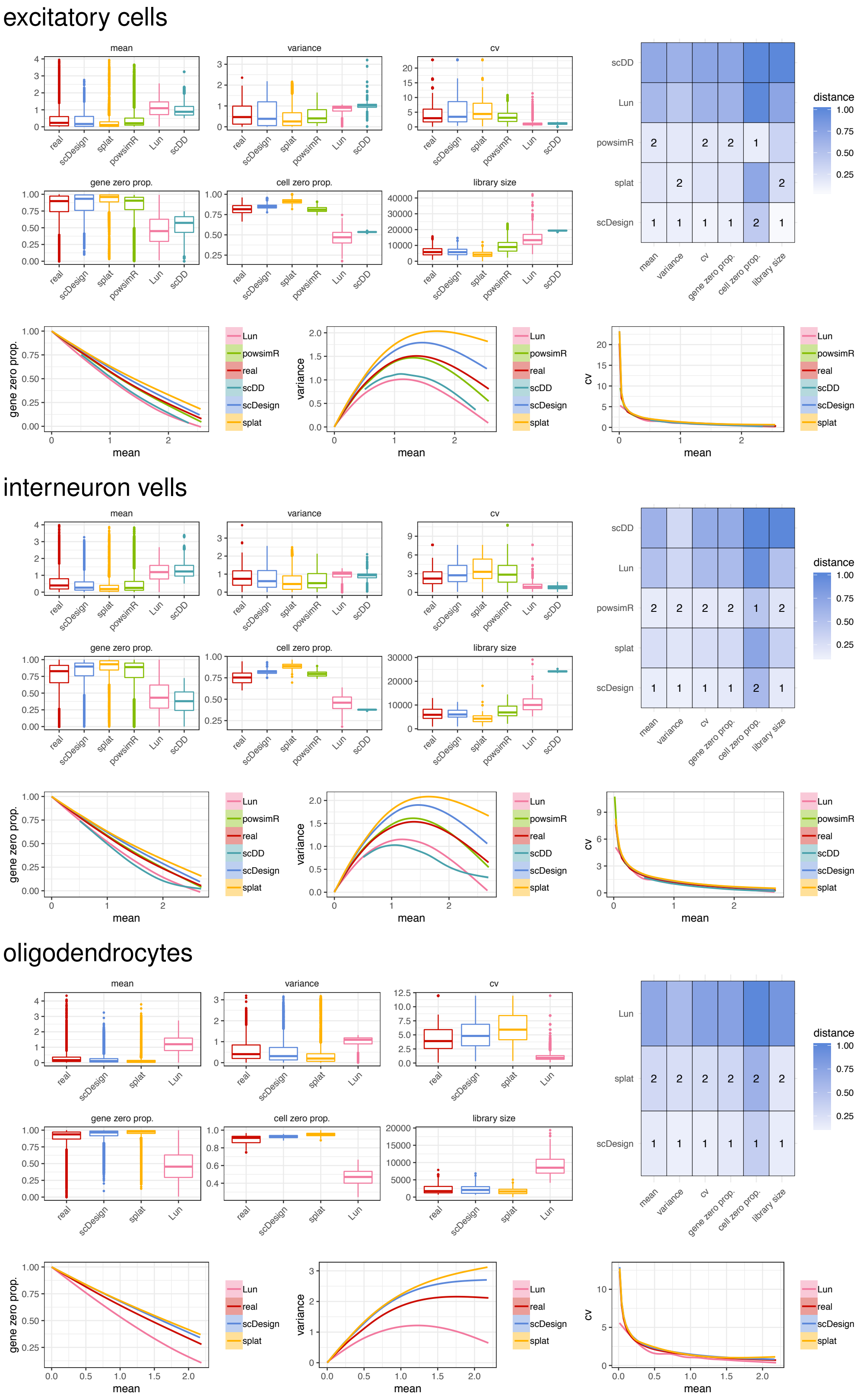


**Figure S7:** Comparison of scDesign and the other four simulation methods based on the inDrop protocol. The boxplots display the gene-wise expression mean, expression variance, expression coefficient of variation, zero proportion, and the cell-wise zero proportion and library size in both real and simulated datasets. The heatmaps display the KS distances between the six statistics in the real data and in the simulated data. The best and second best simulation methods with respect to each statistic are respectively marked with 1 and 2 in the heatmaps. Note that scDD and powsimR failed to simulate data for the oligodendrocyte dataset.

#### Dendrocyte subtype1 vs. Dendrocyte subtype 2

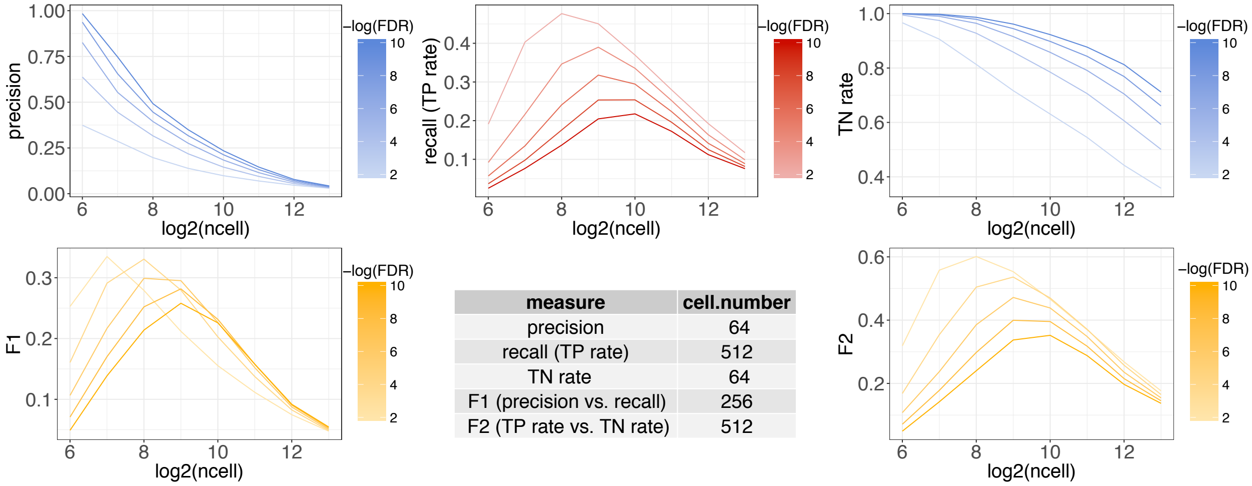


| measure | cell.number |
|---|---|
| precision | 64 |
| recall (TP rate) | 512 |
| TN rate | 64 |
| F1 (precision vs. recall) | 256 |
| F2 (TP rate vs. TN rate) | 512 |

#### Dendrocyte subtype1 vs. Monocyte subtype 1

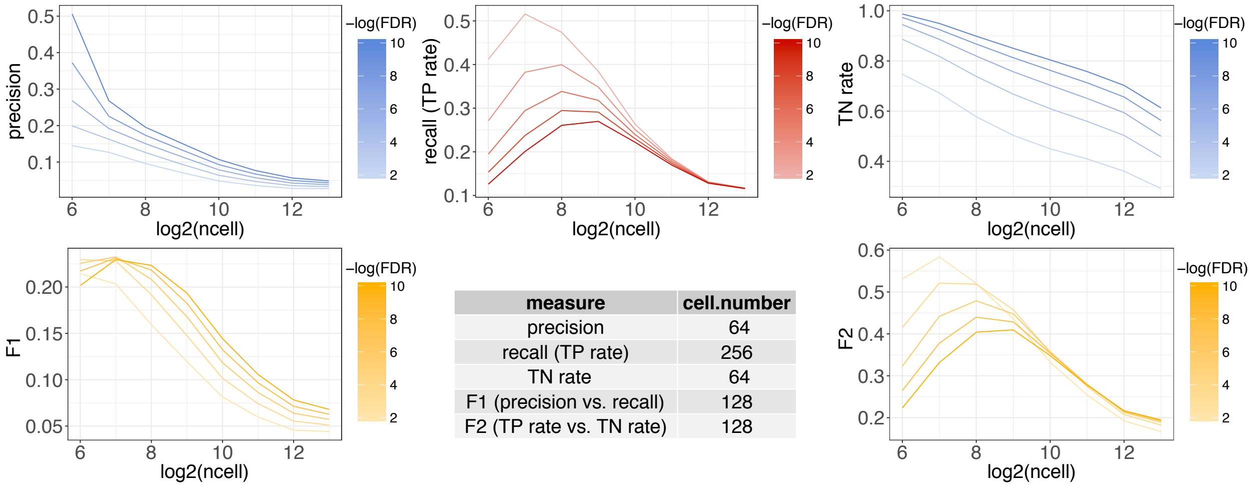


| measure | cell.number |
|---|---|
| precision | 64 |
| recall (TP rate) | 256 |
| TN rate | 64 |
| F1 (precision vs. recall) | 128 |
| F2 (TP rate vs. TN rate) | 128 |

**Figure S8:** Power analysis for DE studies comparing dendrocytes and monocytes (scenario 1). The thresholds on the false discovery rates (FDRs) (to identify DE genes) are denoted in the color legends.

#### CD4 vs. B cells

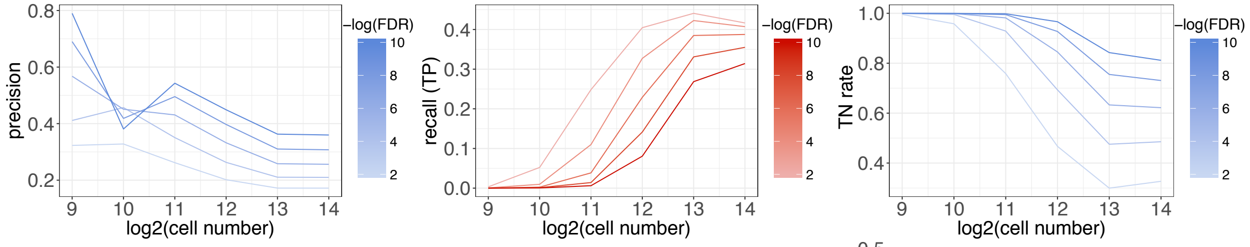


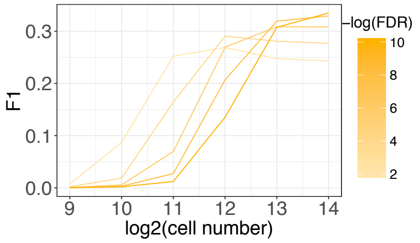


| metric | cell.number |
|---|---|
| precision | 512 |
| recall (TP rate) | 16384 |
| TN rate | 512 |
| F1 (precision vs. recall) | 4096 |
| F2 (TP vs. TN) | 8192 |

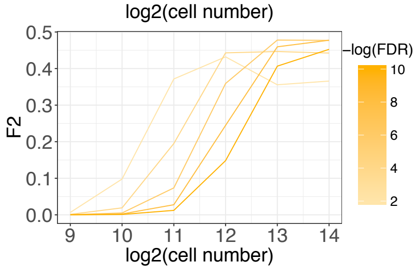


#### CD4 vs. CD8 cells

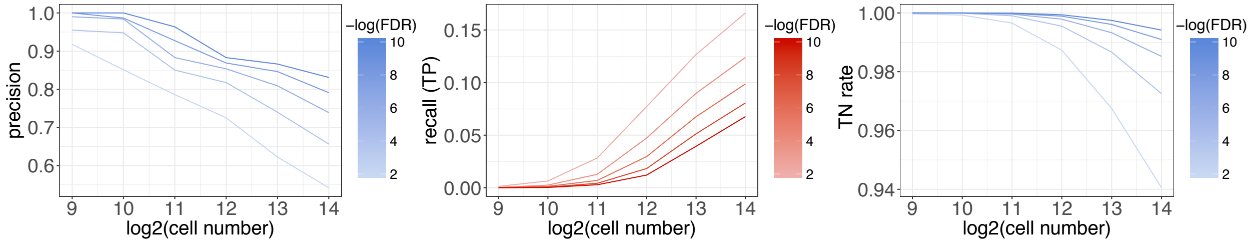


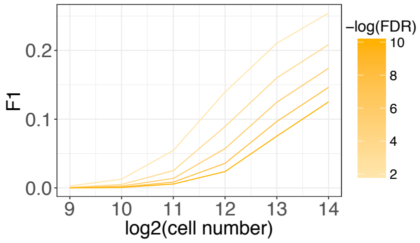


| metric | cell.number |
|---|---|
| precision | 512 |
| recall (TP rate) | 16384 |
| TN rate | 512 |
| F1 (precision vs. recall) | 16384 |
| F2 (TP vs. TN) | 16384 |

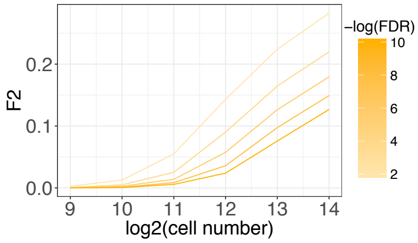


**Figure S9:** Power analysis for DE studies comparing immune cells with the Seqwell protocol (scenario 2). The thresholds on the false discovery rates (FDRs) (to identify DE genes) are denoted in the color legends.

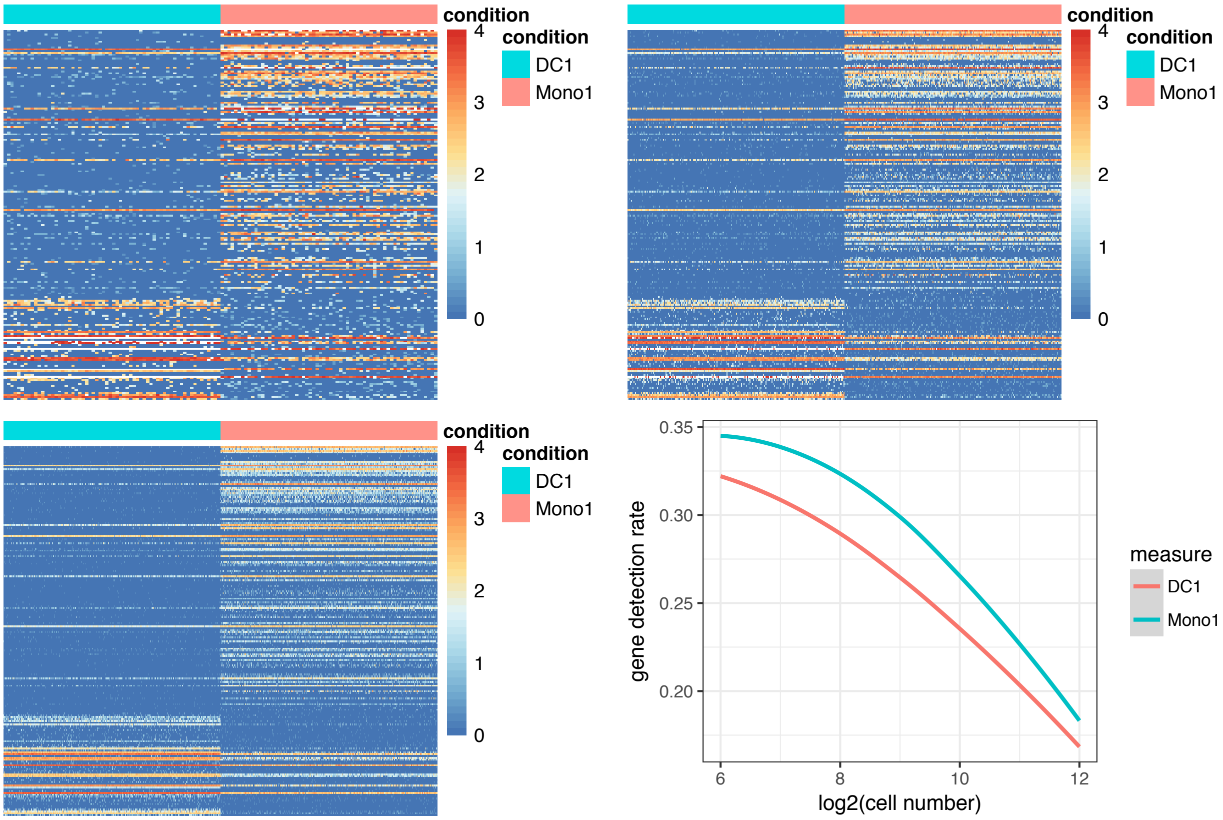


**Figure S10:** Trade-off between cell number and library size given a fixed total sequencing depth. The three heatmaps give the gene expression profiles of the top $200$ DE genes when the simulated cell number of both dendrocytes (DC1) and monocytes (Mono1) is $64$, $256$, and $1024$, respectively. The line plots give the median gene detection rates (i.e., the proportion of genes with non-zero read counts in a cell) in the two cell types as simulated cell number increases.

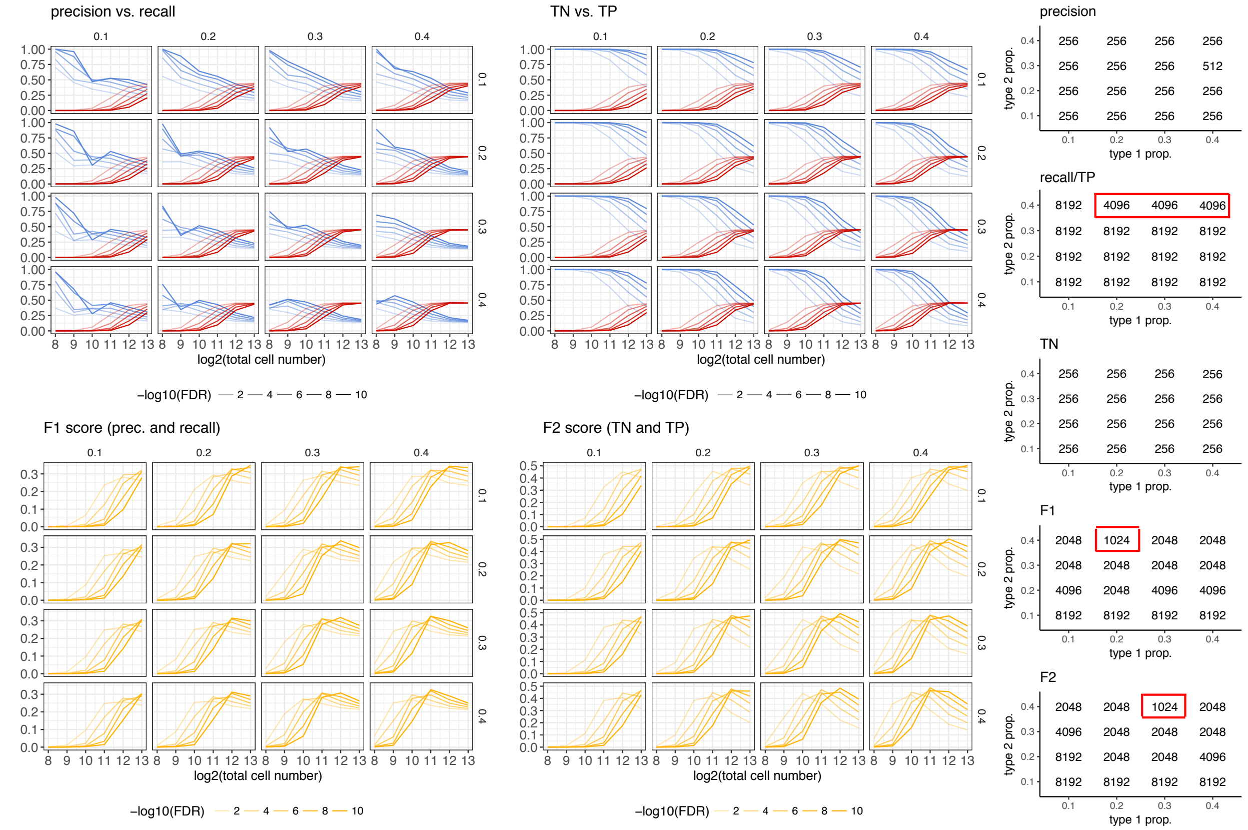


**Figure S11:** Power analysis for DE studies comparing CD4 and B cells with the Seqwell protocol (scenario 2). The thresholds on the false discovery rates (FDRs) (to identify DE genes) are denoted in the color legends. Different proportions ($0.1, 0.2, 0.3,$ and $0.4$) of CD4 and B cells are considered in the experimental design. For the three metrics of recall, F1, and F2, the smallest total cell numbers leading to the best DE accuracy are marked in red boxes.

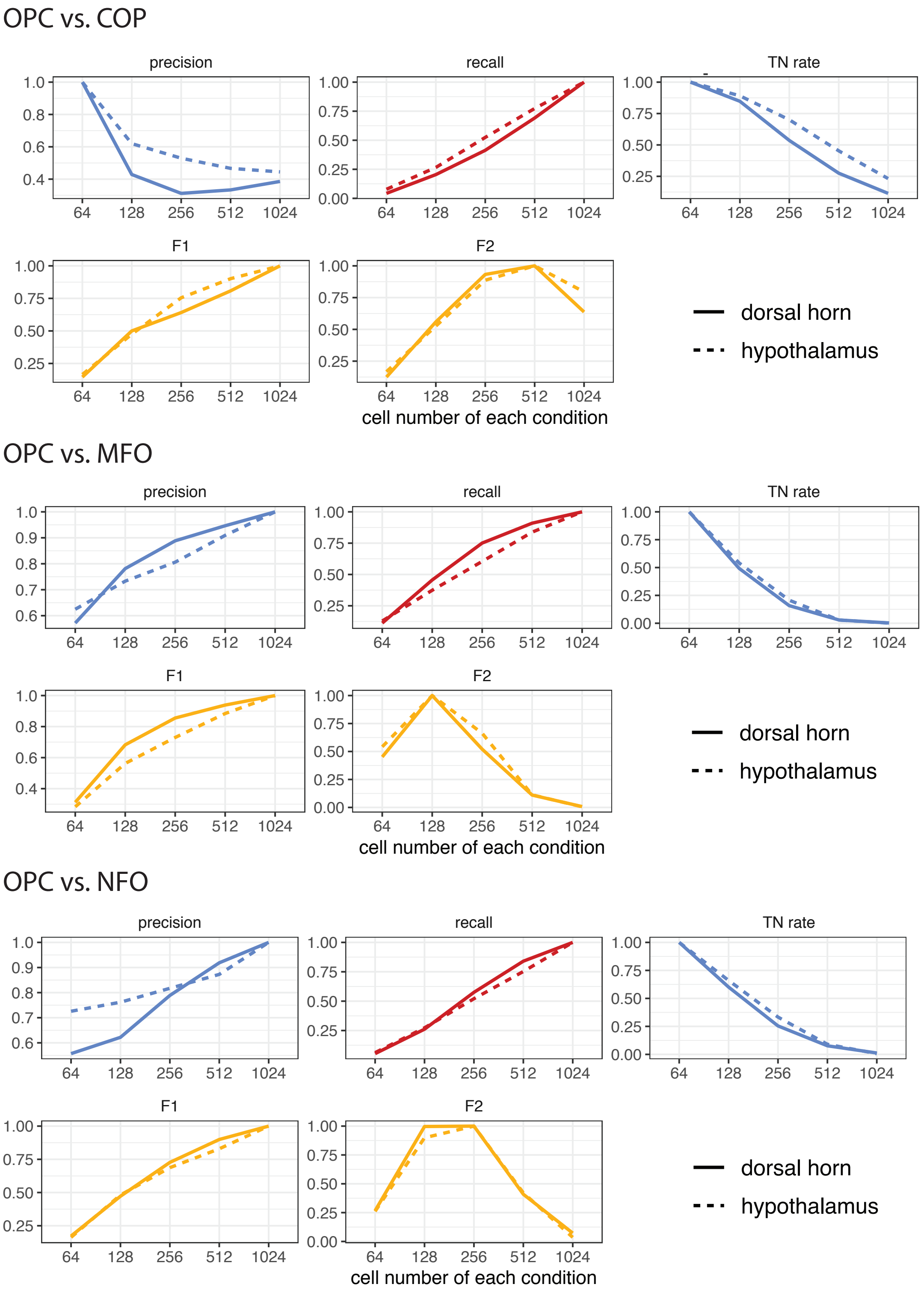


**Figure S12:** Reproducibility of scDesign based on data from different brain regions. The DE studies compare OPC and three other cell types based on scRNA-seq data from two brain regions: dorsal horn and hypothalamus. When identifying the DE genes, the threshold set on the FDR rate is $10^{-10}$. The $y$-axis of each line are divided by the maximum value of that line for normalization.

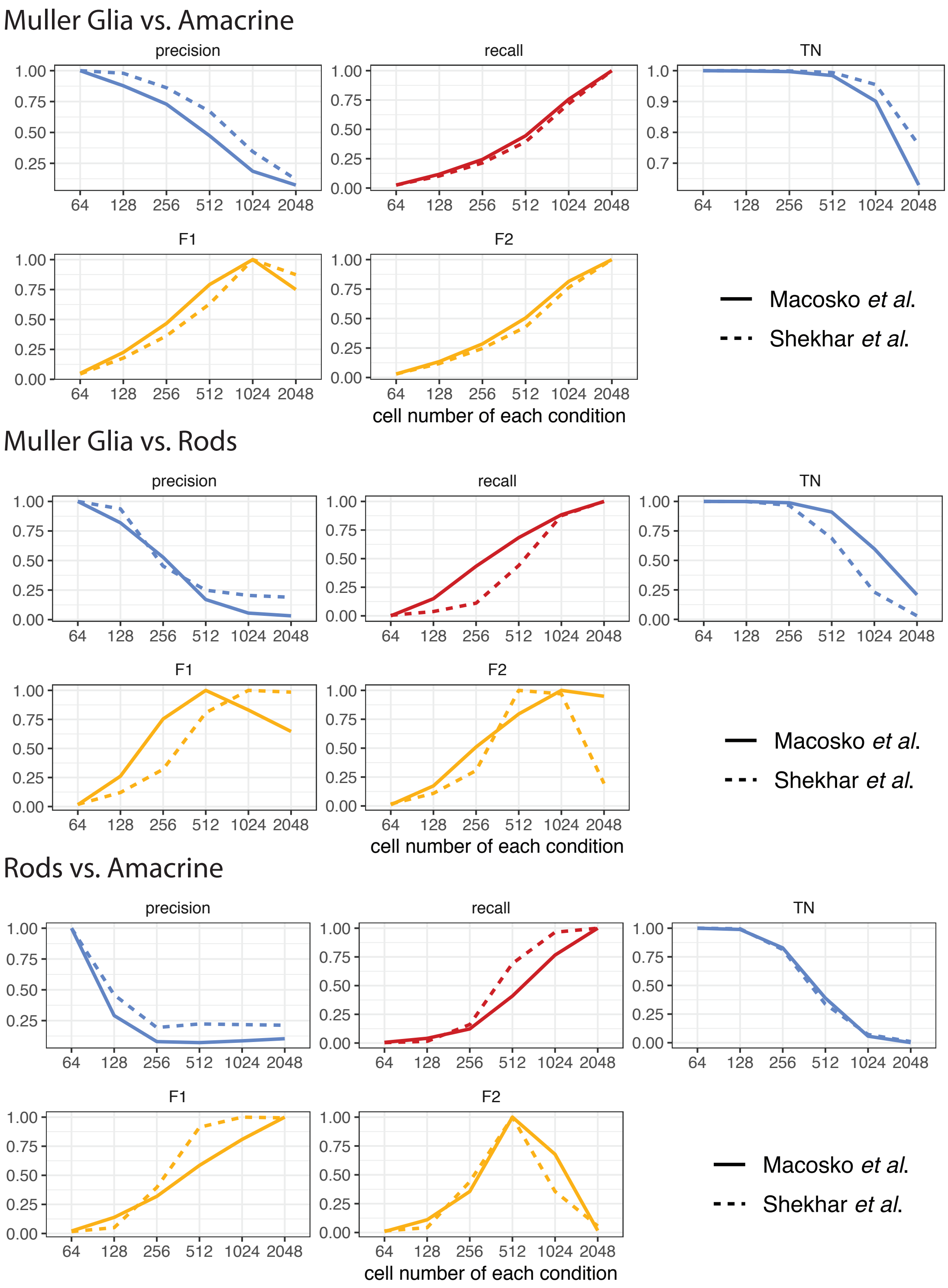


**Figure S13:** Reproducibility of scDesign based on data from different studies. The DE studies compare three types of retina cells based on scRNA-seq data from two studies: Macosko *et al.* and Shekhar *et al.* When identifying the DE genes, the threshold set on the FDR rate is $10^{-10}$. The $y$-axis of each line are divided by the maximum value of that line for normalization.

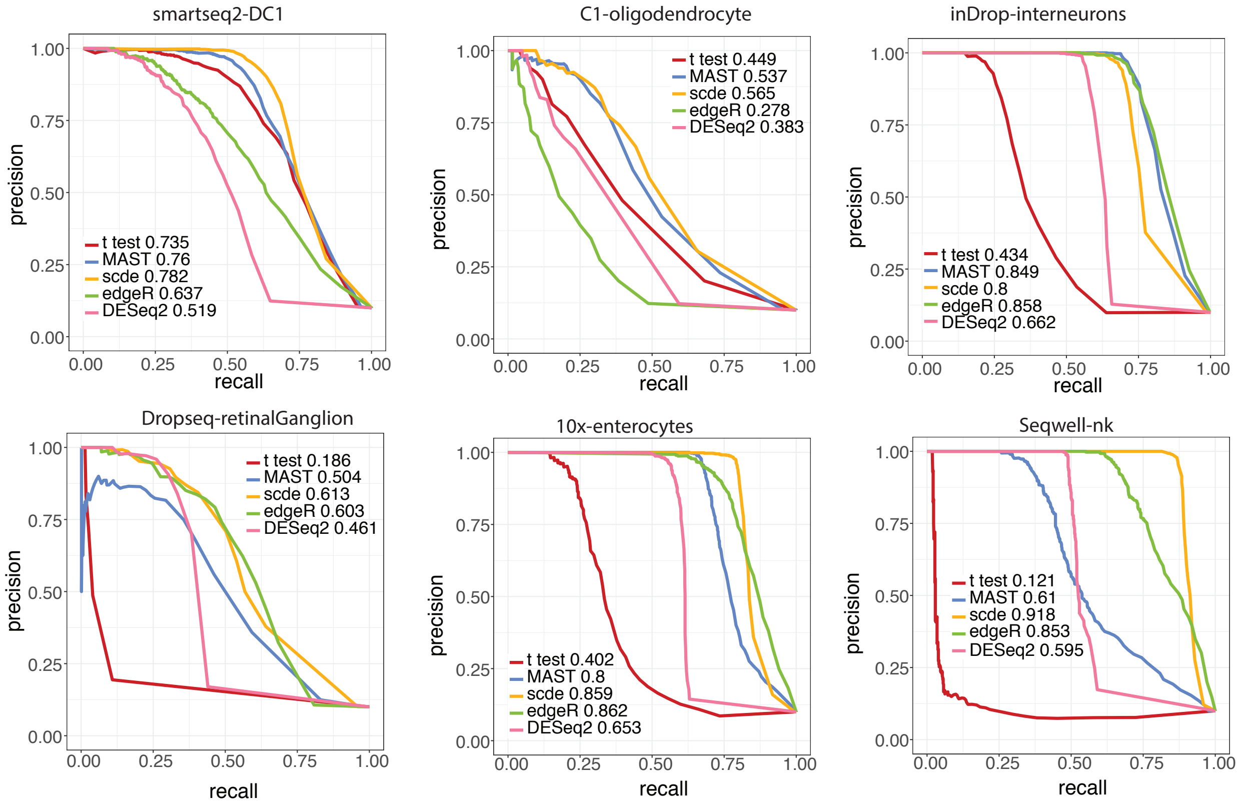


**Figure S14:** Comparison of scRNA-seq DE methods in the first setting. The precision-recall curves of the five DE methods are drawn for the six scRNA-seq protocols, respectively. Corresponding area under the curve (AUC) are given in the plots.

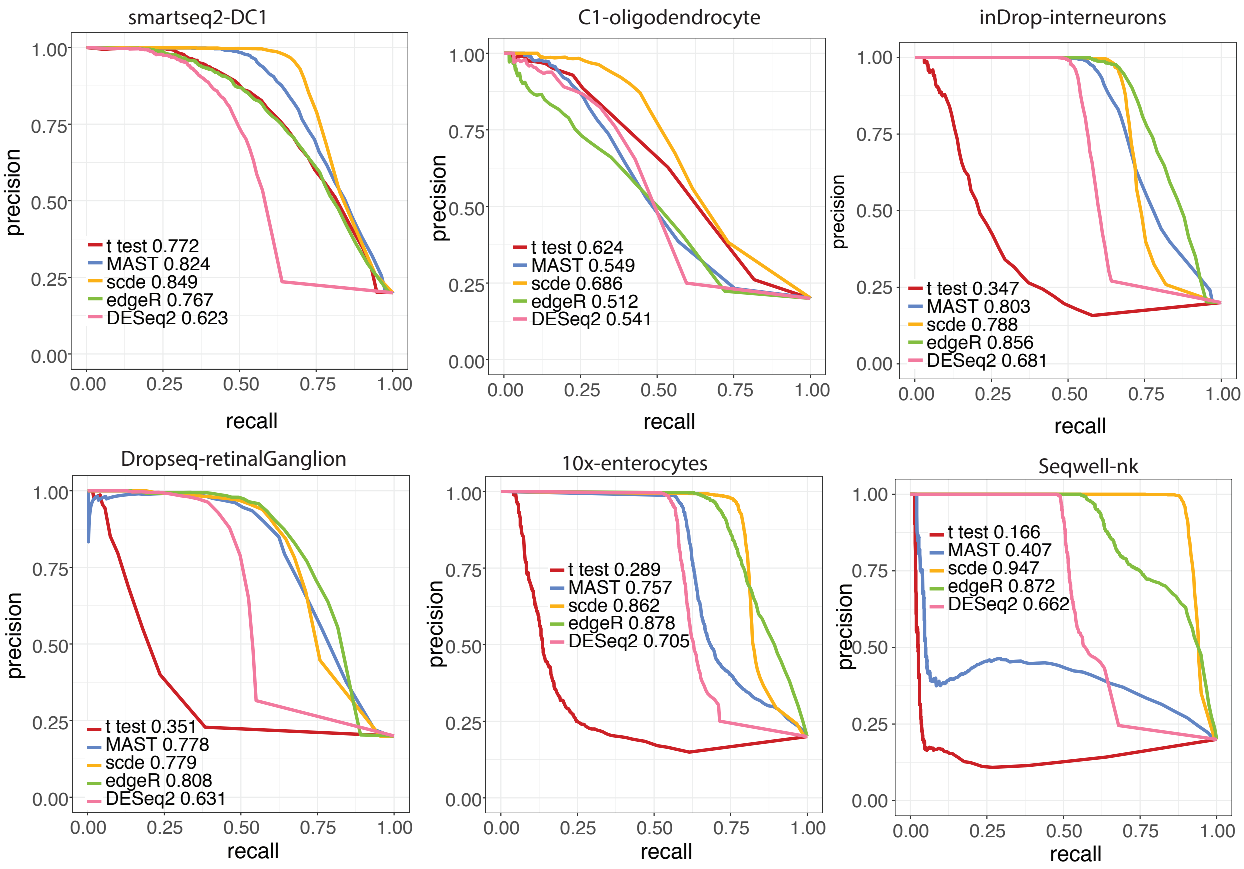


**Figure S15:** Comparison of scRNA-seq DE methods in the second setting. The precision-recall curves of the five DE methods are drawn for the six scRNA-seq protocols, respectively. Corresponding area under the curve (AUC) are given in the plots.

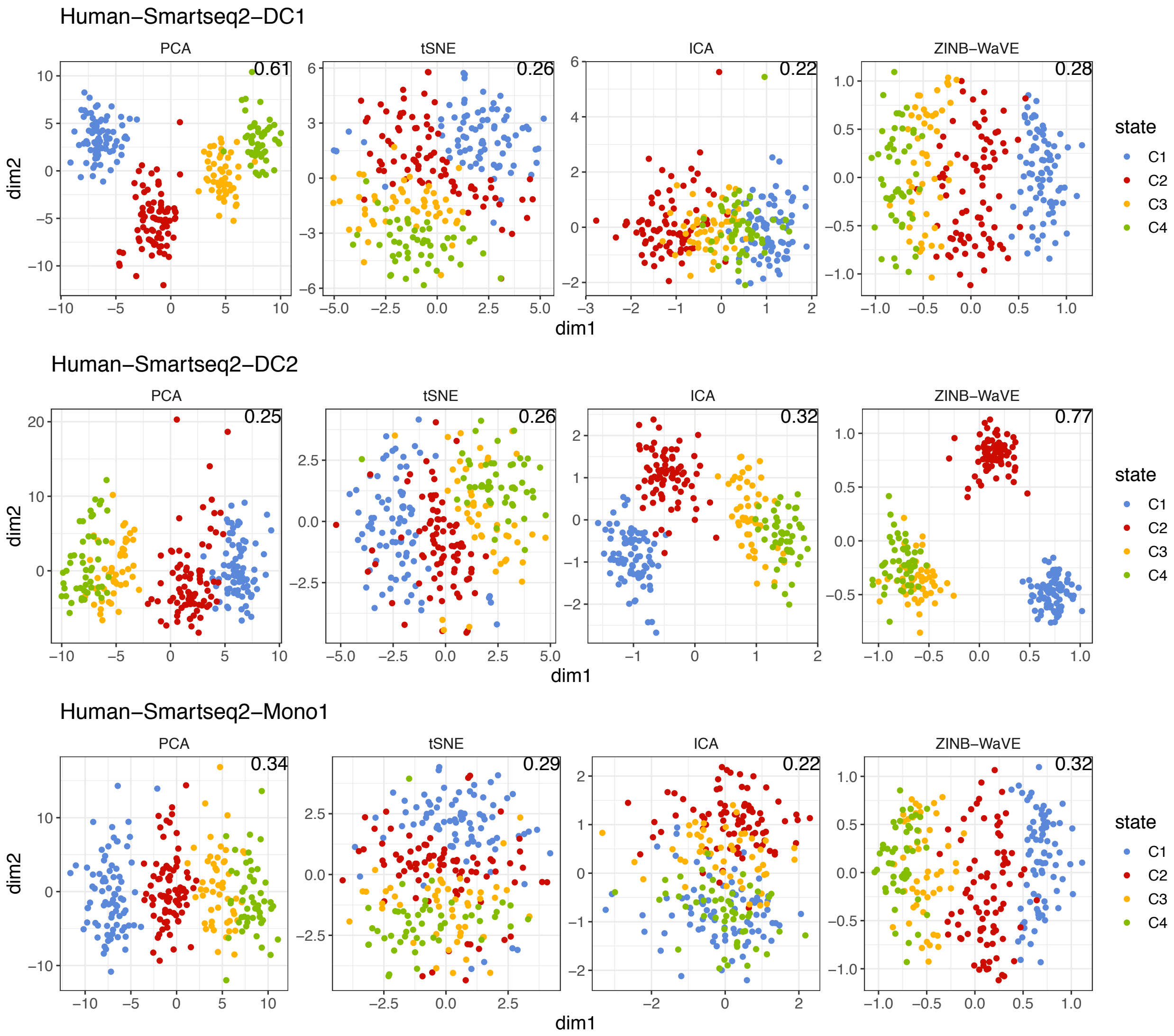


**Figure S16:** Comparison of dimension reduction methods based on the Smart-Seq2 protocol. The first two dimensions resulting from the four dimension reduction methods are given for the data simulated from three cell types, respectively. The hierarchical clustering method was applied on the first two dimensions, and the Jaccard index between the computed cell classes and the true cell conditions is labelled on the top-right of each panel.

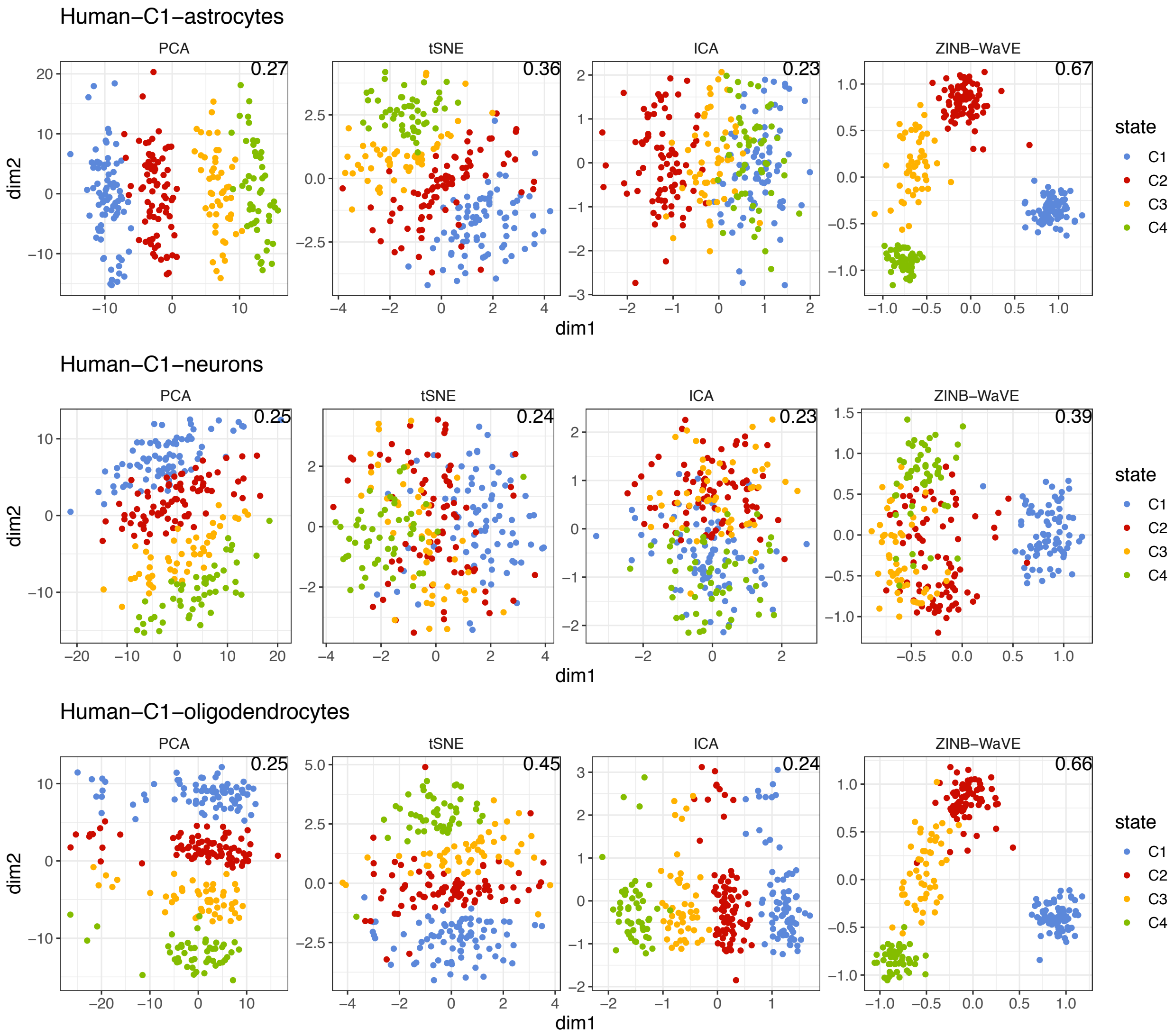


**Figure S17:** Comparison of dimension reduction methods based on the Fluidigm C1 platform. The first two dimensions resulting from the four dimension reduction methods are given for the data simulated from three cell types, respectively. The hierarchical clustering method was applied on the first two dimensions, and the Jaccard index between the computed cell classes and the true cell conditions is labelled on the top-right of each panel.

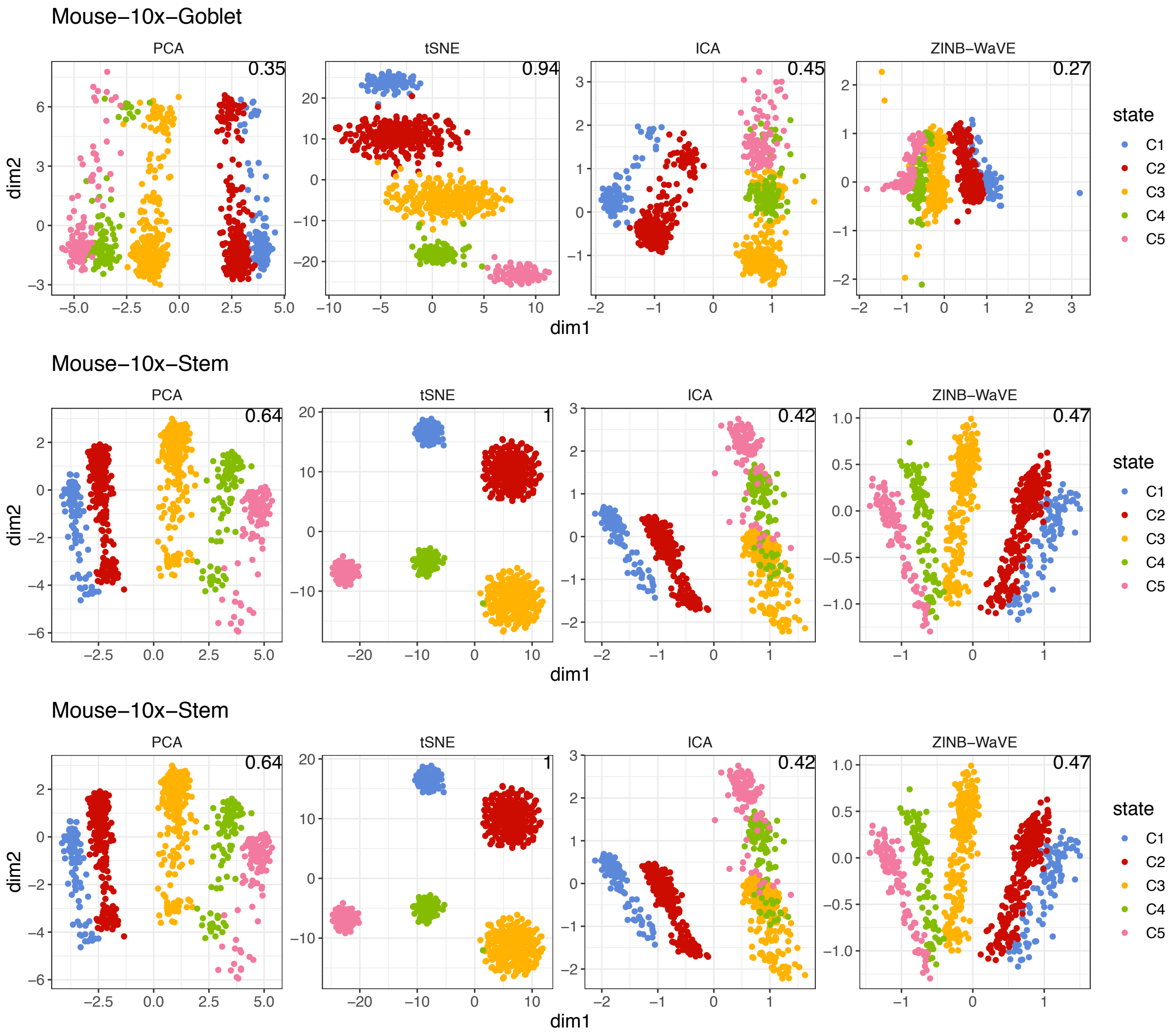


**Figure S18:** Comparison of dimension reduction methods based on the 10x Genomics platform. The first two dimensions resulting from the four dimension reduction methods are given for the data simulated from three cell types, respectively. The hierarchical clustering method was applied on the first two dimensions, and the Jaccard index between the computed cell classes and the true cell conditions is labelled on the top-right of each panel.

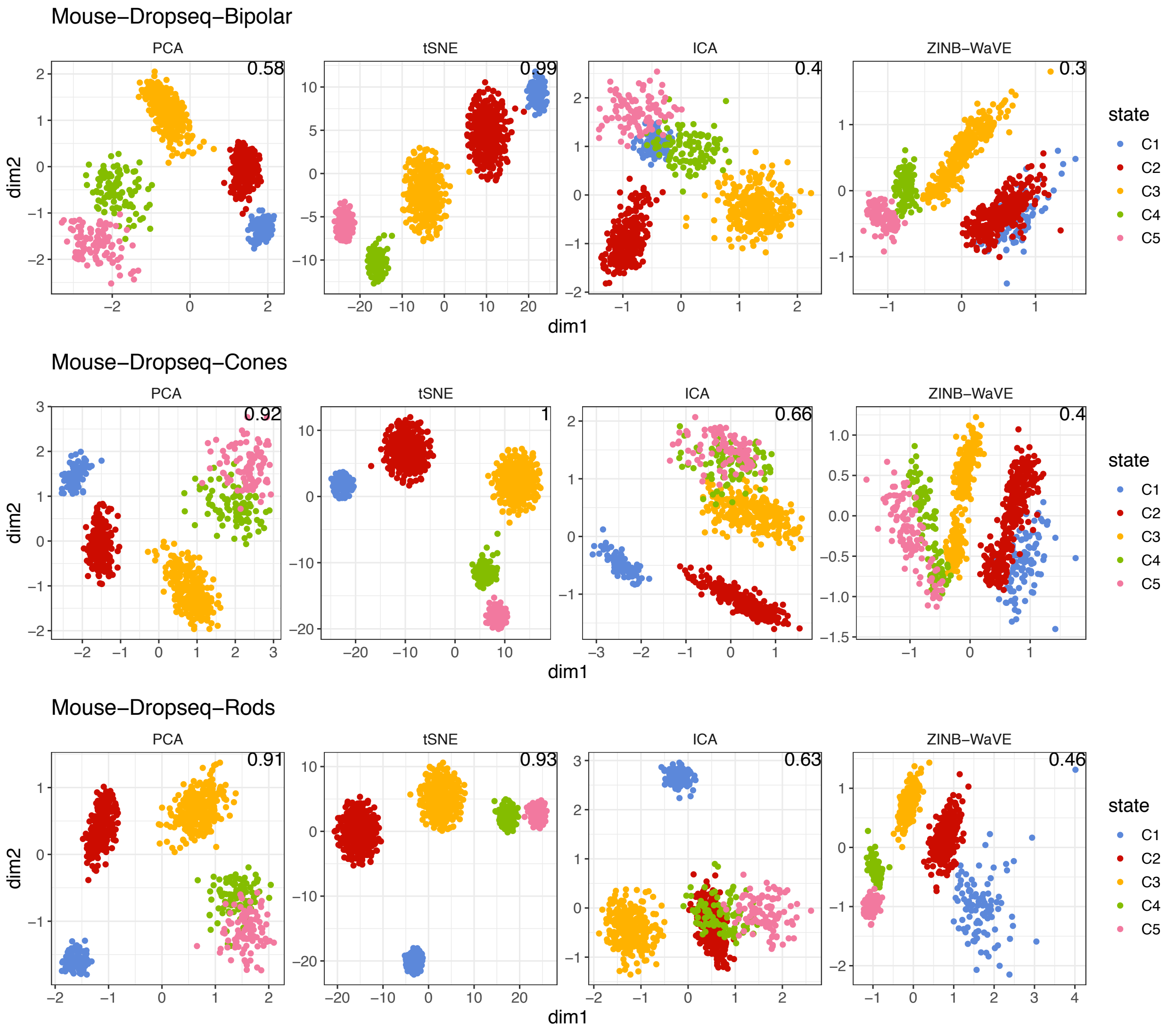


**Figure S19:** Comparison of dimension reduction methods based on the Drop-Seq protocol. The first two dimensions resulting from the four dimension reduction methods are given for the data simulated from three cell types, respectively. The hierarchical clustering method was applied on the first two dimensions, and the Jaccard index between the computed cell classes and the true cell conditions is labelled on the top-right of each panel.

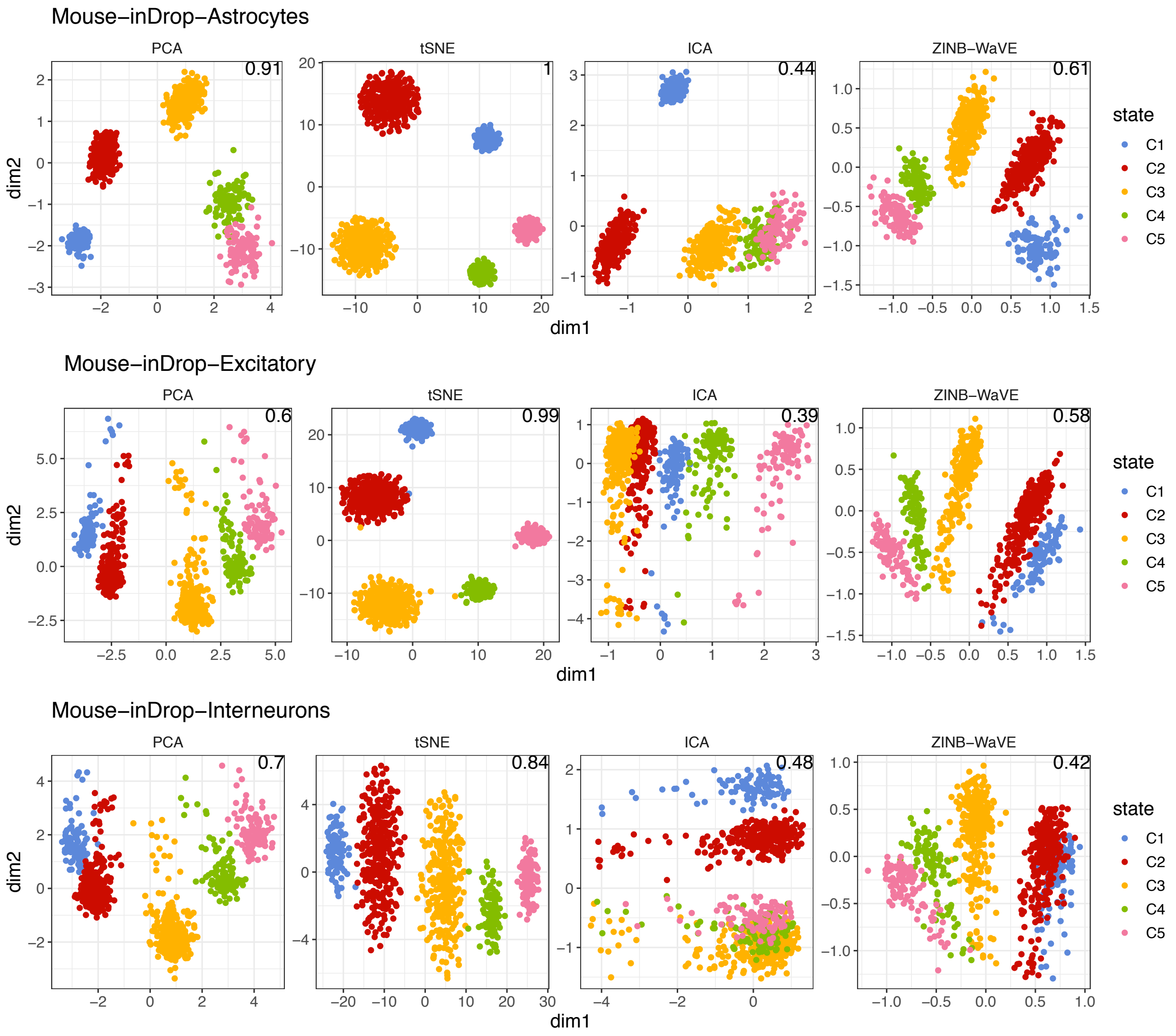


**Figure S20:** Comparison of dimension reduction methods based on the inDrop protocol. The first two dimensions resulting from the four dimension reduction methods are given for the data simulated from three cell types, respectively. The hierarchical clustering method was applied on the first two dimensions, and the Jaccard index between the computed cell classes and the true cell conditions is labelled on the top-right of each panel.

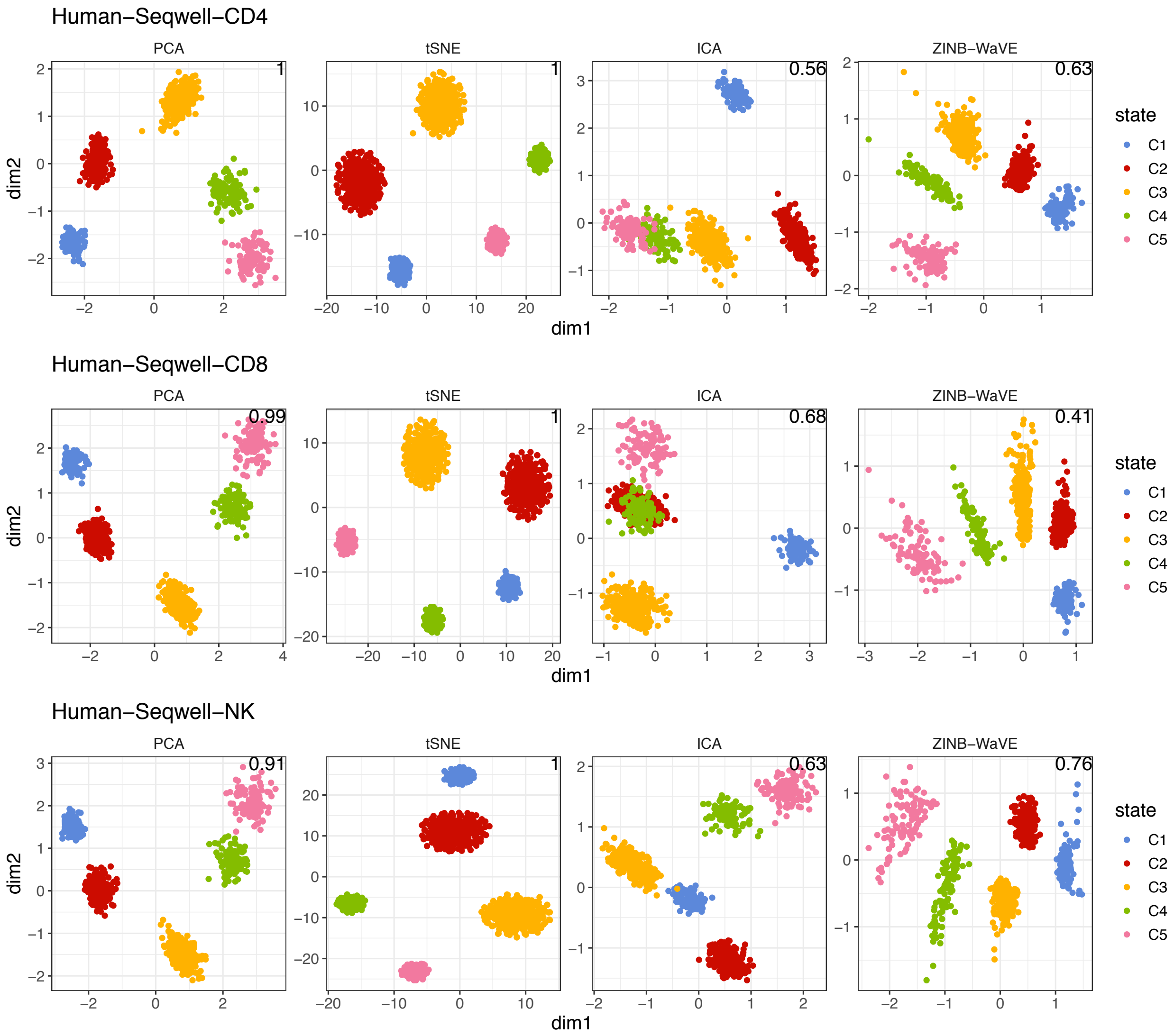


**Figure S21:** Comparison of dimension reduction methods based on the Seq-Well protocol. The first two dimensions resulting from the four dimension reduction methods are given for the data simulated from three cell types, respectively. The hierarchical clustering method was applied on the first two dimensions, and the Jaccard index between the computed cell classes and the true cell conditions is labelled on the top-right of each panel.
